# Cell shapes decode molecular phenotypes in image-based spatial proteomics

**DOI:** 10.1101/2025.05.13.653868

**Authors:** Trang Le, William D. Leineweber, Matheus P. Viana, Anthony Cesnik, Jan N. Hansen, Wei Ouyang, Susanne M. Rafelski, Emma Lundberg

## Abstract

The diversity of cellular and tissue structures can arise from a few basic cell shapes, which undergo various transformations based on biophysical constraints on cytoskeletal organization. While cellular geometry has been linked with selected biological processes such as polarity, signaling or morphogenesis, the orchestration of the whole proteome in association to cell shape is still poorly understood. In this study, using more than 1 million images of single cells stained for 11,998 proteins across 10 cell lines in the Human Protein Atlas database, we performed an integrated analysis of organelle, pathway and single protein levels in association to a 2D cellular shapespace. We found that cell and nuclear shapes across cell lines exist in a shared continuum. We also found that the subcellular organelle topology varies across cell lines, but remains robust within each cell line’s shapespace. At the single protein level, we found that cells of different shapes in the same cell cycle phase might be preparing for different fates, and that many non-cell cycle proteins expressed shape-based abundance variation. Using the same coordinate framework defined by shape, we could analyze the distribution shift of protein spatial localization under drug perturbation.

## Introduction

Shape has been a useful descriptor of cellular phenotypes ever since Robert Hooke coined the term “cell” in 1665^1^. Hundreds of years later, cell shape remains critical for identifying cells, owing to the principle that form and function are tightly associated. The shape of a cell is the complex output of a large network of molecular interactions, meaning that the cellular shape should be able to provide information on its molecular state. This relationship has been applied across biology and medicine. Pathologists use cell morphology to diagnose diseases^2^. Researchers also rely on cell shape to differentiate cell types and functions, cancerous from non-cancerous cells^3^, stem cell differentiation^4^ or epithelial-to-mesenchymal transitions (EMT) and downstream metastasis^5^. Cell shape, particularly plasma membrane curvature and eccentricity, has also been mathematically identified to be a locus of retrievable information storage in cells^6^. However, all these studies were conducted comparing shapes in a control versus perturbed condition. The natural range of shape variation and the associated protein expression and spatial distribution remains elusive. A recent study on the variation of 3D shapes of an induced pluripotent stem cell (iPSC) population found that the intracellular topology of organelles is robust across shape variation, and that for specific events, like early mitosis, the reorganization of organelle topology (*i.e.* change in the association between subcellular organelles) is associated with the rewiring of the cell needed to support the cell division process^7^. These findings reinforce the importance of studying what constitutes the normal range of expression and localization of subcellular components, and subsequent cell state definitions.

To understand the molecular underpinnings of various cell shapes, it is essential to study proteins as the key structural and functional molecules in the cell. Proteins are hierarchically organized into functional units^8^ and subcellular protein localization not only dictates specific cellular processes but also provides critical insights into mechanisms, such as cell division^9,10^, nucleus-cytoskeleton reorganization in cellular migration^11,12^, wound healing^13^ and tissue folding^14,15^. For example, translocation of Cyclin B1 from the cytosol to the nucleus is a key checkpoint in the cell cycle^16^. These intricate spatial subcellular patterns of proteins can be studied with molecular imaging. Fluorescence microscopy has led to extensive imaging databases that cover nearly all intracellular proteins across numerous cell types (*e.g.,* Human Protein Atlas or HPA^17^, OpenCell^18^). The HPA has compiled images of nearly all intracellular proteins (13,534 protein-encoding genes, v24) across multiple cell types. However, information about the subcellular localization of each protein was collected independently, with each sample containing a single protein staining and three reference markers. While it is possible to co-visualize dozens to hundreds of proteins at a subcellular level using multiplexed staining, such as with 4i^19^ or CODEX^20^, these techniques are currently limited to around hundred proteins at best. Consequently, there is a need to develop a unified approach for systematically studying how protein subcellular organization correlates with overall cell morphology, such as a landmark-based common coordinate framework, which has been used for other spatial data types (*e.g.,* brain MRI scan, or mouse brain gene expression)^21,22^.

There have been many recent efforts to move beyond qualitative descriptors of cell shape into more powerful and quantitative approaches to measure cell shapes ^6,7,23–25^. Most work to date has focused on understanding organelle function and distribution in relation to cell shape^7^, commonly using a mode of quantification highly optimized for the organelle in question, such as blob detection for vesicles and intensity-based thresholding for more continuous structures such as ER sheets and actin filaments^26^, while some studies developed more generalized approaches for all organelles^7^. While cell shape parameterization is not new, extracting meaningful biological insights from shape-based models remains difficult due to the limited integration of molecular data, as most studies focus on well-characterized proteins within established systems^27^. However, the proliferation of tools and approaches for quantifying cell shape and protein distribution speaks to the recognized potential of cell shape as a high-throughput readout of molecular states. Despite these advances, a standardized framework to explicitly map all organelles structures to the cell shape continuum, a common coordinate framework defined by cellular shapes, remains a critical challenge. Such a framework is essential for scaling proteome-shape studies from large public databases and identifying new targets for downstream hypothesis-driven studies.

Given the ability to associate image-based proteomic data with shapes, a number of outstanding questions can be addressed to systematically advance cell shape analysis: What are the common shape variations within a cell line? What can cell shape tell us about its intracellular organization of organelles and proteins, and how are they distributed with respect to each other and across cell types? Do cell morphologies relate to specific cell states (*e.g.,* metabolic states), or can multiple phenotypes manifest in shapes indistinguishable from each other? How are these (re)organizations linked to cell state and function?

In this paper, we seek to characterize nuclear and cellular shape variations within and across cell types and map proteome-wide data to this shape-based common coordinate framework. Leveraging the HPA^17^, we determine the relationships between cell shape and organelles across cell lines, known cell states and cell processes as well as expression and localization of individual proteins. Our analysis reveals that the principal modes of cell shape variation are conserved across diverse cell types. Subcellular organelle organization is shape-invariant within cell lines but different across cell lines. Although shape alone is a suboptimal predictor for interphase progression, many cell cycle dependent proteins show similar intensity trajectories related to nuclear shape. We also found that cells displaying different shapes in G2 might be preparing for different cell fates based on their metabolic proteome profiles. We demonstrated that comparing shape-matched cell subpopulations instead of whole populations in perturbation data allowed us to reveal drug effects on protein expression that are obscured in studies on the whole population level. Furthermore, we observe protein localization shifts following certain drug treatments, such as a protein that is normally distributed across multiple cellular compartments in untreated cells becomes uniformly localized to the nucleus. This may indicate that the drug is arresting cells in a specific state.

## Results

### Robust determination of a cellular shapespace across cell lines

To explore shape variations and the corresponding proteome profiles within and across populations of cells, we selected ten cell lines from the HPA v23^17^ with diverse morphologies, including carcinoma, sarcoma, non-cancerous epithelia, and myeloid-derived cell lines (Fig. 1A). A total of 862,111 individual cells were segmented to determine the nuclear and cellular shapes, including between 5,242 to 262,802 individual cells from each cell line (Supplementary Table 1). Observing heterogeneous shapes within and across cell lines prompted us to develop a comprehensive pipeline aimed at identifying the principal modes of shape variation^7^ (Fig. 1B). Fourier transforms of cell and nucleus shapes were analyzed using principal component analysis (PCA) to determine the major axes of variation. Inverse Fourier transform enabled visualization of an “average” cell shape at the center point of the principal components (PCs). Moving along each PC while keeping the others at center, we can observe the corresponding shape changes exemplified by each axis, which we termed “shapemodes” (Fig. 1C, Supplementary Table 2).

**Figure 1.**
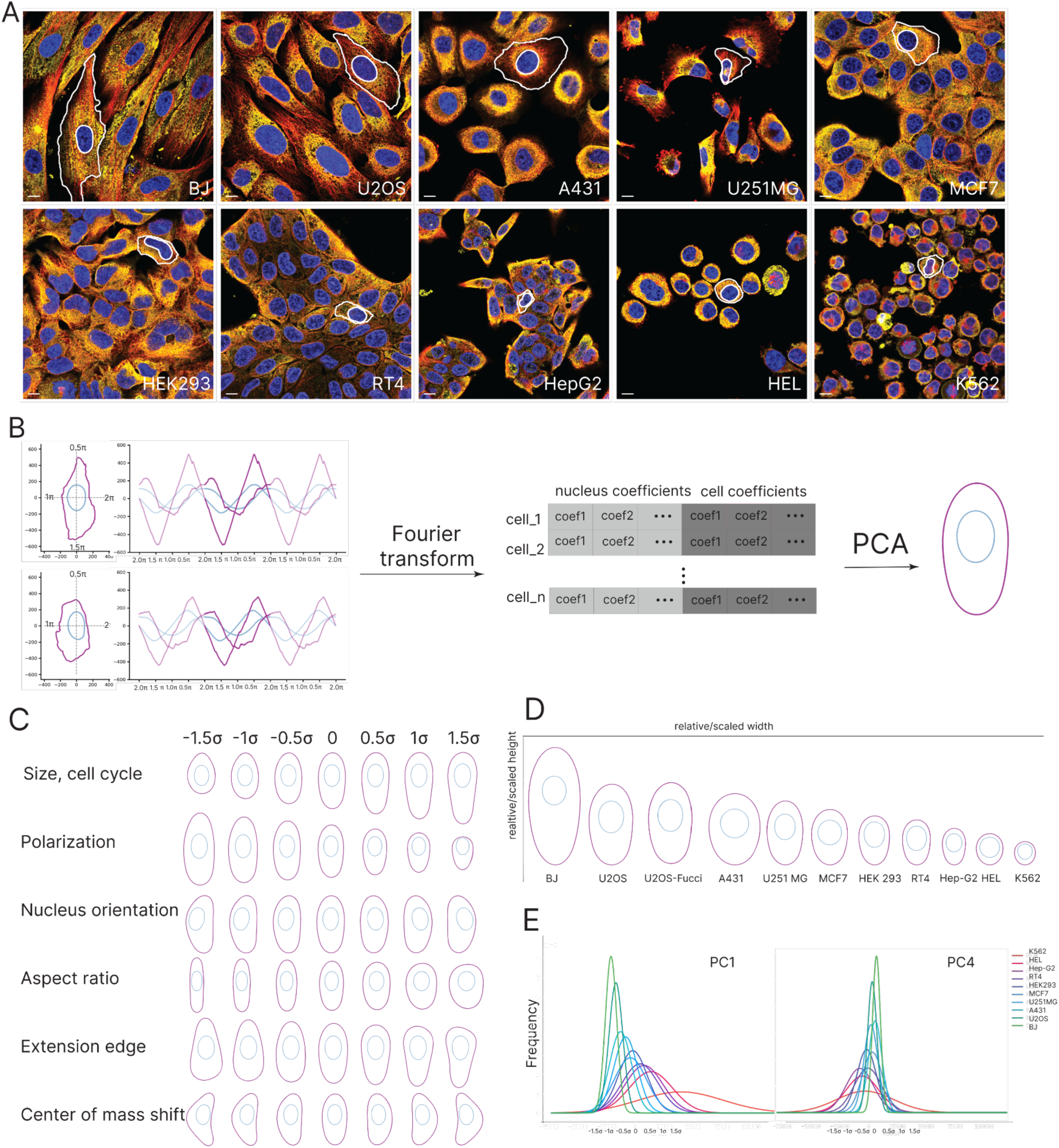
Cell line morphology and reconstruction of the cell and nucleus shape space. (A) Representative morphology of 10 distinct cell lines, with white outline of nuclear and cell shapes. Scale bar 10 μm. (B) Schematic of the shapespace construction pipeline: raw images are processed to extract individual cell and nuclear shapes, followed by Fourier Transform (FFT)-based shape coefficient extraction. These nuclear and cell shape coefficients are reduced to main modes of variations (shapemodes) by Principal Component Analysis. Average cell shape could be reversed transformed from the shapespace. (C) U-2 OS shapespace consists of different shapemodes (variation along each principal component while freezing the other components), which account for ∼ %90 variation of the shape coefficient matrix for each cell line. (D) Mean cell shape representations across the 10 cell lines, capturing characteristic morphological differences such as shape, size, aspect ratio, etc. as well as average representation of U-2 OS cells under different imaging objectives and conditions (U-2 OS dataset was captured at 63x with a confocal, and U2OS-FUCCI is captured at 40x with a widefield microscope, average cell of U2OS-FUCCI was rescaled to displayed the same average nucleus and cell shapes for the same cell line). (E) Distribution of cell types within shapemodes representing PC1 and PC4 of the common shapespace, illustrating shared and distinct morphological features among cell lines.

The first six PCs, accounting for ∼90% of the variance for each cell line, collectively represent the observed “shapespace” of a measured cell population, and could be well described by human-interpretable shape descriptors (Fig. 1C, Supplementary Figure 3). These shapemodes were nearly identical across cell lines (Supplementary Figure S1), with the first two PCs (representing cell size and polarization) accounting for the greatest variance in cell shapes (Supplementary Figure S1). The first shapemode (PC1), which accounted for 38-53% of the total variance across datasets (Supplementary Table 2), reflected changes in cell size and polarization while maintaining nuclear consistency. Conversely, variations along PC2 corresponded to nucleus elongation along with the cell shape (variance explained ∼ 16-22%). These shapemodes are robust across different imaging modalities for the same U-2 OS cell line (Supplementary Figure S2). The similarity of the shapespace across diverse cell types and imaging modalities indicates that our method to parametrize cell shape is robust and that cell shape heterogeneity may arise from conserved underlying processes.

We also observed that the cell line–specific shapemodes closely matched those from the common shapespace—constructed by randomly sampling cells from all lines—underscoring the robustness of our approach. However we could observe sight variations in the distributions, for example myeloid-derived cell lines, such as K-561 and HEL, displayed tight distributions along PC1 (size, elongation) and PC4 (aspect ratio), whereas mesenchymal-derived cells, such as BJ and U-2 OS, had broader distributions (Fig. 1E). Despite the differences in distributions between cell lines, they occupied a continuous shapespace rather than being clustered to distinct regions of the common shapespace. This finding further indicated that variations in shapes represent a continuum both within and across cell lines.

### Cell lines have distinct organelle topologies

Having established that nuclear and cellular shape variations are robust across cell lines, we sought to investigate organelle organization changes across cell types and shapemodes. To perform this analysis, we quantified 5 to 18 organelles (for each cell line) by aggregating all single cells that were stained for proteins associated uniquely with a specific organelle (i.e. no multilocalizing proteins) as determined using a localization classification model developed on the HPA images^28^. Unlike previous studies that relied on a single representative marker per organelle^7^, this approach aggregated the stainings of multiple (dozens to hundreds) proteins per organelle (Supplementary Table 3), thereby enhancing the robustness and accuracy of our organelle topology assessment (Fig. 2A,B).

**Figure 2.**
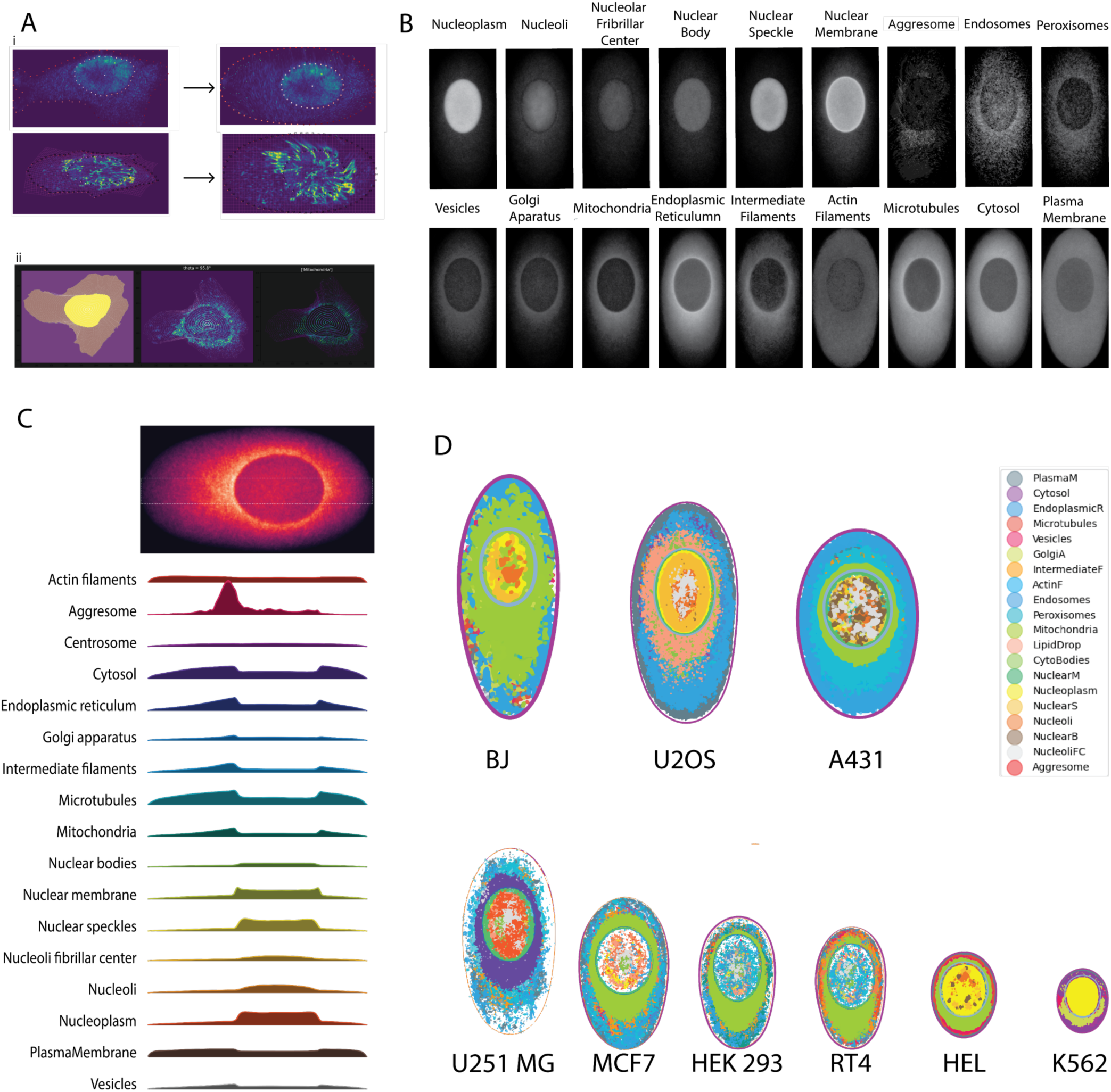
Protein spatial parameterization and average organelle representation of up to 20 organelles across 10 cell lines. (A) Two approaches for single-protein spatial representation: (i) Morphing: using a thin plate spline deformation field computed from cellular landmarks (nuclear centroid, nuclear segmentation, and cell border segmentation) to transform original representation to a continuous protein intensity map in the average cell shape; (ii) Sampling: defining equidistant sampling points in concentric rings extending from the nuclear centroid to the nuclear boundary and then to the cell periphery, with protein intensity values sampled with Gaussian kernel. (B) Spatial distribution of organelles visualized as the average organelle representation, exemplified in U2OS cells. Each organelle map was constructed by aggregating the binarized morphed single cell samples that express the specific organelle pattern, and whose cell and nuclei shapes were at the middle (std=0) shape bin. (C) Organelle density distribution along the cell’s longest axis, revealing spatial trends. (D) Average organelle representation mapped onto the mean cell shape for each of the 10 cell lines, illustrating cell line-specific organelle organization.

Using the integrated intensity distribution for each organelle across different cell lines, we procured an average organelle representation per shapemode and cell line which facilitated comparative analyses of how organelle localization varies by cell shape (Fig. 2B). Specifically, single cell protein localization was parameterized into intensity profiles by morphing according to deforming fields calculated from landmarks (nucleus centroid, nucleus contour and cell contour) (Fig. 2 A&B). This approach overcame limitations of sampling at concentric rings^7^, which can lead to over- or under-parameterization in the cytosol. Aggregation of all proteins in the average shape can be visualized as spatial organelle probability maps within a given cell shape (Fig. 2C). With such maps we revealed, for example, that aggresome distribution is highly polarized, with the highest probability of observing aggresomes in the perinuclear region on the elongated side of the cytosol. Overall average representation maps of organelles present in 10 cell lines are visualized in Fig. 2D.

Depending on the cell line and data availability, we were able to assess the colocalization of 5 (in the average K-562 cell shape) to 18 cell organelles (in the average U-2 OS cell shape, shown in Fig 3A-J) using the determined average organelle representations. Correlation analysis revealed two clusters of organelles whose spatial maps were highly correlated in all analyzed cell lines, corresponding to the metacompartments of sub-nuclear and sub-cytosolic organelles (Fig. 3). By contrast, representation maps of endosomes, vesicles, and the ER varied most across cell lines. This could indicate that vesicles and the ER may be either flexibly positioned across cell types or tie to characteristic cell functionalities. For example, the ER occasionally demonstrated a positive correlation with the nucleus, indicating localization above or below the nucleus, as a result from 2D imaging. Such flexibility appears intuitive for vesicles, which are known to dynamically move within the cell. Yet, the observed variability may also reflect fundamental differences in the average cell shapes across different cell lines, which appear to align with the differences in organelle topology (Supplementary Figure S3).

**Figure 3.**
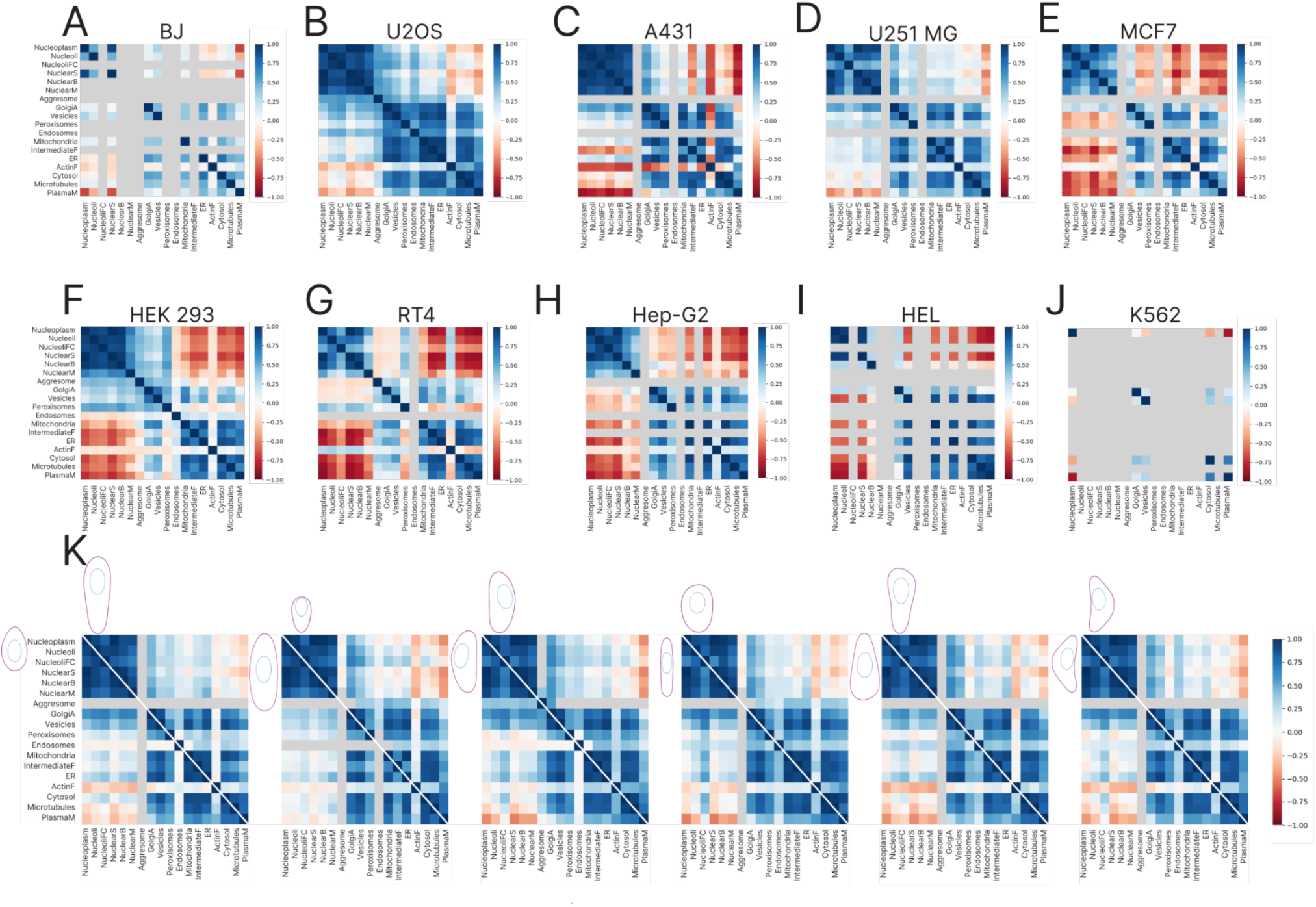
Organelle Correlation Across and Within 10 Cell Lines. (A–J) Organelle spatial representation colocalization patterns exhibit distinct spatial relationships across different cell lines while maintaining robustness within individual cell lines. Representation for each organelle in each cell line was constructed by different and overlapping sets of proteins associated with each cell line highlighting key biological pathways that can be further explored using the corresponding stained gene sets. (See Supplementary Figure S6 for additional details). Heatmaps were used to show correlations between data-supported average organelle representation maps present in each cell line, relationships from -1 to 1, gray means not enough sample to have a representative map. (K) Organelle organizations within cell lines are consistent despite morphological variability (displayed by 2 extreme groups for each shapemode in U2OS shapespace).

### Organelle associations are robust across shapespace in the same cell line

We further investigated the organelle topology within the same cell line across shapes to test the hypothesis that the reorganization of major cellular components correspond to shape variations. Using a similar approach as that used to analyze the organelle topology across cell lines, we found that the spatial distribution of organelles proved to be robust across cell shapes (heatmap inconsistency score ∼ 0.01 ± std) (example in Fig. 3K). This consistency suggests a conserved subcellular organization within cell types despite shape variability.

### Highly variable protein variation through shape changes support key biological processes

Having established that organellar topologies are robust within cell lines, we examined the expression of individual proteins within these organelles. In a normal log-phase growing cell population, most protein expression levels remain stable; however, about 26% (3,275 out of 12,502) of proteins exhibit single-cell variation in intensity or spatial distribution based on manual annotation (HPA v23)^29^. To explore whether these variable proteins correlate with cell shape, we analyzed a dataset of 1,160 highly variable proteins visualized in U-2 OS cells genetically modified with the FUCCI reporter system^29,30^. We first trained a segmentation model, enabling the measurement of cell shape, and protein abundance (i.e. staining intensity) in both the nucleus and cytoplasm. We then analyzed these highly variable proteins using pathway assessments, employing grouped t-tests, Kruskal-Wallis tests and permutation analysis to determine if protein concentrations in either compartment vary across principal components.

At the pathway level, we found that the 1,160 highly variable proteins were enriched for 11 biological pathways from the Human Molecular Signatures Database (Enrichr adjusted p-value < 0.05; Fig. 4A, Supplementary Table 4). Testing protein variability along shapemodes revealed that 633 proteins vary across different shape projections (number of proteins with FDR < 0.05: 374 for PC1, 143 for PC2, 523 for PC3, 182 for PC4, 92 for PC5, 87 for PC6, Supplementary Table 5). We found that two shapemodes relating to cell size (PC1 and PC3 for FUCCI U-2 OS cells) exhibited the most significant protein variations in shapespace. When mapping these proteins to their associated pathways, we observed a strong enrichment in cell cycle and growth pathways. For instance, the Mitotic Spindle pathway contained 5 significantly variable proteins out of 26 matched proteins, and the G2-M checkpoints pathway had 9/40 proteins significantly variable through PC3. Pathways associated with cell growth, such as KRAS^31^ and mTORC1^32^, also exhibited a high proportion of significant protein associations with size and nuclear polarity related shapemodes. Beyond the size and nuclear polarity, PC4, which differentiates elongated versus round cells, showed a high proportion of significantly variable proteins involved in metabolic pathways, including Glycolysis, Fatty Acid Metabolism, and Xenobiotic Metabolism (Fig. 4A). In addition, we also found that a number of proteins showed variations along shapemodes independently of the cell cycle, including understudied proteins^33^ such as C11orf96, C3orf18, C10orf82, SPATA12, C9orf85, HHIPL2, KIAA1143, FAM241A. These results suggest that distinct cellular shapemodes can encode biologically meaningful variations at pathway and protein levels.

**Figure 4.**
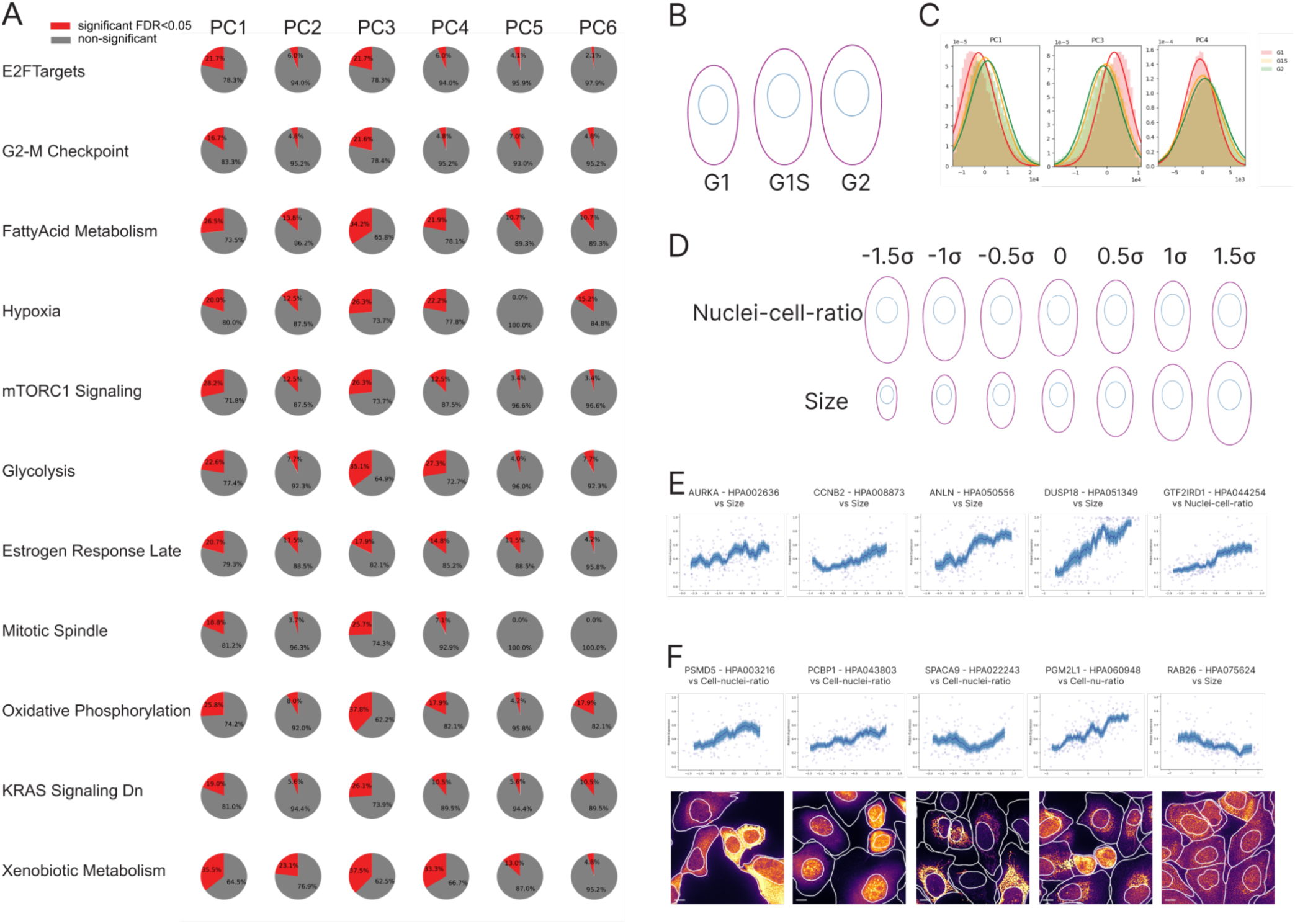
Cell Cycle dissection on shapespace. (A) Pathway enrichment analysis of the dataset, highlighting the proportion of proteins that show significant variation in shapespace. (B) Average cell shape representations of cells in G1, G2S, and G2 cell cycle phase, illustrating expected cell and nuclear changes as the cell cycle progresses. (C) Original shapemodes with the highest distribution shift. (D) Independent component analysis (ICA) of the same shape coefficient matrix resulted in two robust shapemodes that best capture the cell cycle signal when orthogonality constraints are removed. (E) Example of cell-cycle-dependent proteins exhibiting expression trajectories that align with shape changes throughout the cell cycle. (F) Example of non cell-cycle-dependent proteins that show shape-associated expression variations, accompanied by representative sample images. Panels E–F displayed compartment-specific quantification of single-cell protein expression, emphasizing spatial regulation at the subcellular level. Scale bar 10 μm.

### Shapemode dissection of the cell cycle

Cell cycle progression is accompanied by drastic changes in nuclear and cellular size and shape. The shapemodes of cells in G1, G1/S and G2 phases are identical to each other and to the shapemodes of the entire cell population (Supplementary Figure S2), indicating that cell shapes in different cell cycle phases follow common axes of variation. Cell and nuclear size displayed an expected increase in G2 compared to G1S compared to G1 population, as well as a shift in the shape distribution towards more rounded cells (Figure 4B). Based on nuclear and cell contours alone, it is challenging to determine the cell cycle phase (3-class classification accuracy similar to using nuclear size alone, ∼0.55, see Methods). We observed a large distribution overlap between cell cycle phases (Fig. 4C). Thus, even though changes in average DNA content, nucleus size and cell size reportedly can reveal cell cycle characteristics of a whole cell population^34–36^, we conclude that cell and nucleus shape distributions alone are not robust classifiers of the cell cycle interphase position in U2OS cells.

Using independent shape components (ICA, see Methods), we identified two shapemodes that scale with cell cycle progression (Figure 4D). Over one fifth of cell-cycle dependent (CCD) proteins (20.6%, 73 of 354) displayed changes in protein abundance correlated to changes in the shapemodes. Examples include classic cell cycle checkpoint proteins, such as AURKA^37^, CCNB2, ANLN^38^, and other CCD proteins such as DUSP18, GTF2IRD1, KIF23 ^29^ that varied with cell and nuclear size or cell-to-nucleus ratio (Fig. 4E, Supplementary Table 6). Additionally, we observed proteins not previously linked to the cell cycle that vary with these two ICA-identified shapemodes (permutation test, see Methods). For example PSMD5, PCBP1, SPACA9 and PGM2L1 show changes in expression correlated to the cell to nuclei ratio, while RAB26 exhibits an overall decreasing trend with increasing cell size (Fig. 4F) .

To determine whether cells of the same cell cycle phase but different shapes exhibit distinct molecular profiles, we compared the metabolic pathways of G2 cells with large round versus large elongated morphologies (Fig. 5A). We found that large round cells have higher purine and pyrimidine metabolism, oxidative phosphorylation, while large elongated cells have higher sphingolipid metabolism (Fig. 5B). Upregulated expression of proteins involved in nucleotide metabolism like purine^39^ (AK6, XDH in Fig. 5C, MPO etc) and pyrimidine (UPP1, GPT2 in Fig. 5D) might indicate that these G2 large round cells are preparing for DNA replication and division. Increase in multiple proteins of Complex I (NADH:ubiquinone oxidoreductase) in the mitochondrial oxidative phosphorylation system (NDUFA1, NDUFA9, NDUFV1 etc) might indicate increased ATP production, aiding the cell division process. Conversely, the increase in enzymes involved in sphingolipid metabolism, including lipid-related proteins like LPIN2, OGT, TMPPE (Fig. 5E) may indicate increased membrane rigidity^40^, which helps maintain the membrane tension needed in elongated shaped cells. These results indicate that even within the G2 cell population, rounder cells are more metabolically active preparing for division, while another group of elongated cells may be focused on other processes such as migration.

**Figure 5.**
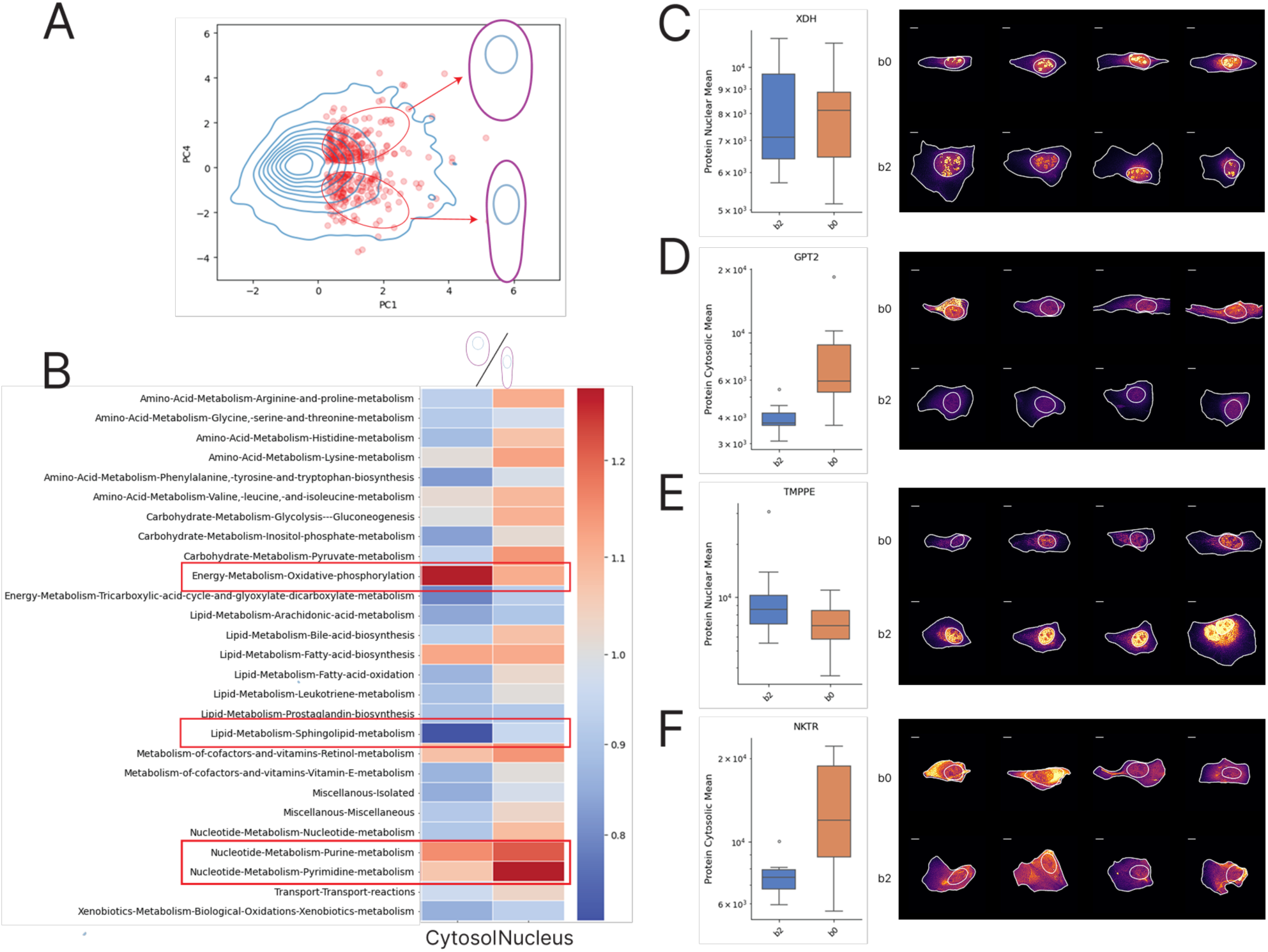
Differential metabolic profiles of G2 cells on different shape groups. (A) Cell shape distribution (blue kernel density contours) of 20,000 randomly sampled cells, highlighting two distinct G2 subgroups (red sample points): round & large G2 cells and elongated & large G2 cells for comparison. (B) Fold change analysis of metabolic pathways between the 2 cell populations. (C-F) Examples of proteins that show significant differences in expression level between elongated and round cells. Left panels: Bar plots of log fold changes in protein abundance between the two shape groups. Right panels: Representative cell images from each group (top row: elongated cells; bottom row: round cells), Scale bar 10 μm. (C) XDH in Oxidative phosphorylation pathway. (D) GPT2 in Purine/pyrimidine metabolism pathway. (E) TMPPE in Sphingolipid metabolism pathway. (F) NKTR in Isolated enzyme pathway.

### Protein variation under perturbation through the lens of the shapespace

Understanding the approximate bounds of protein variation within an unperturbed system, and considering that organelle topology remains relatively stable with respect to shape changes, we aimed to investigate whether notable shifts occur in either the morphology or protein expression levels under drug perturbation. For this we used a dataset of more than 30,000 breast cancer cells (cell line MDA-MB-468), treated with paclitaxel, a microtubule stabilizer, vorinostat, a histone deacetylase (HDAC) inhibitor, or untreated. Each cell was stained for cellular reference markers (DNA, microtubules, ER) and one of 95 different proteins related to chromatin remodelling ^41^.

Analysis of the shapespace in cell populations that are untreated and treated with one of the two drugs revealed that PC1, associated with size, and PC4, which differentiates between round and elongated shapes, exhibit the most significant shifts in shapespace distribution. Specifically, cells treated with Paclitaxel (which inhibits microtubule depolymerization and blocks cell division) appeared smaller and rounder compared to normal cells (with a Wasserstein distance of 0.18, compared to 0.03 between untreated cells and Vorinostat) (Fig. 6A).

**Figure 6.**
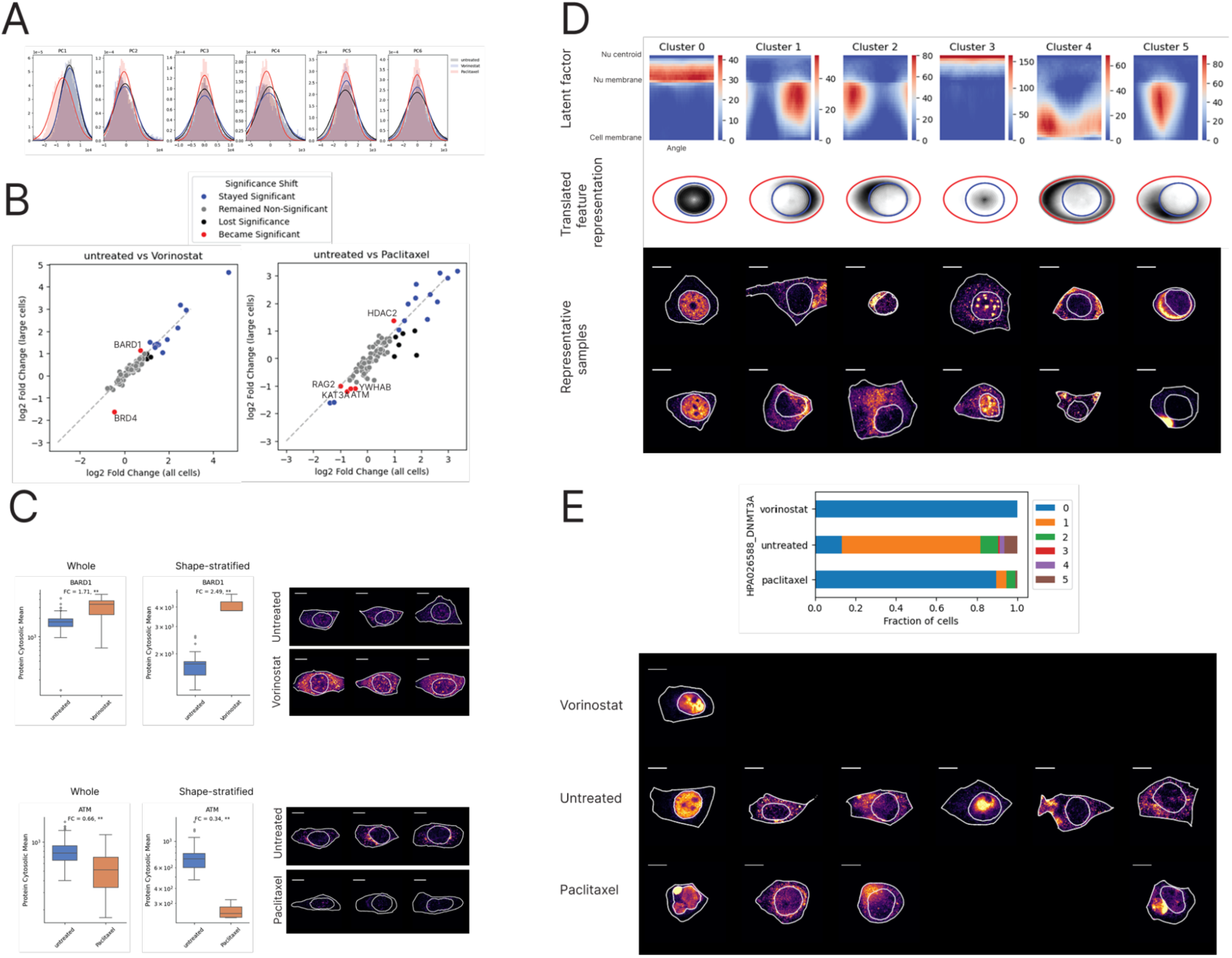
Cell shape and protein distribution shift upon perturbation. (A) Distribution shifts in cell size and elongation axes under perturbed conditions. (B) Increased effect sizes observed in subgroup-based intensity analysis (C) Example of proteins whose expression level was detectably different only with shape-constraint analysis: The proteins encoded by BARD1 or ATM, perturbed with vorinostat or paclitaxel, respectively. (D) Latent factors of protein expression derived from intensity parameterized matrices. The angle axis represents a whole circle. (E) Example protein (the DNA-methyltransferase 3A encoded by the gene DNMT3A) with diverse spatial locations in untreated cells but localizing only to the nucleus upon treatment of vorinostat. Scale bar 10 μm, intensity for single cells linearly rescaled to 99th percentile of all protein-specific images for display.

Importantly, shape-constraint comparisons of untreated and treated cell subpopulations yielded larger effect sizes for detecting changes in protein expression for some proteins. For example, new proteins were found to significantly differ in expression comparing treated and non-treated cells when looking at elongated cells rather than population averages (Fig. 6B). The shape-stratified up/down-regulated hits included the proteins BARD1 (Fig. 6C), ATM (Fig. 6C), PRMT1, MSL1, PRKAA1, RAG2.

Projecting all protein expression patterns into the common coordinate framework with nuclear and cell landmarks, we observe six major protein localization patterns: nuclear (cluster 0), centroid (nucleoli pattern that gets averaged to a midpoint, cluster 3), nuclear periphery on the smaller cytosolic side (cluster 1), cell periphery (cluster 4), and two cytosolic regions on the larger side (cluster 2 and 5) (Figure 6D). With each cell assigned one of these patterns, we compared the distribution of treated cells to untreated cell spatial locations. Several proteins exhibited heterogeneous subcellular localization patterns under unperturbed conditions but converged to a single compartment (usually nucleus) following treatment with vorinostat, such as DNMT3A or TAF1 (Figure 6E, 6F). This phenomenon could be attributed to vorinostat’s specific mechanism of action, which induces cell cycle arrest predominantly at the G2 phase^41^.

## Discussion

Shape is a complex trait that is influenced by many underlying molecular processes. With a systematic association analysis of protein variation across shapespace, we resolved the relationship of subcellular protein localization with nuclear and cellular shape variation, at the level of organelles, pathways and single proteins.

The shapemodes in this paper are similar to those identified in a study on 3D shape and organelle distribution in human iPS cells^7^. This provides external validation of our findings and supports the idea that 2D images, like those used in our study, can capture sufficient insight into cell shapes and organizational structures in many cell lines — reinforcing the conclusion in the referenced iPSC-focused study that organelle association is robust across shapespace. By employing multiple proteins per organelle in our approach, we have strengthened the reliability of the organelle representation across cell types. The preservation of major shapemodes in our study of a 2D image dataset compared to the reference 3D study may result from cell surface tension^4,42,43^ being detectable in 2D images, and that each cell type has an inherent distribution along a shared continuum.

We found that, in unperturbed cells, the association between cell morphology and state is convoluted. Most likely, multiple cell states relate to similar cell shapes indistinguishable from one another. These slight variations in morphology still guarantee similar organelle topology within a cell type, which indicates that there is a stable cell architecture within a population of cells of the same type.

We also found that each cell line has a specific organelle topology. These differences may reflect cell-line characteristic functions, such as related to protein trafficking dynamics, or signaling networks. While expression levels of compartment-specific proteins in each cell line may fluctuate, the cell line-specific spatial relationships among organelles remain stable throughout shapespace, preserving overall cellular architecture and functional integrity. This insight highlights the strong potential for generative or representation deep learning models to learn the underlying distribution of organelles across cell lines, paving the way towards virtual cell foundation models^44^. We also anticipate that more use of machine learning in modeling the interpretable physical forces underlying cell-shape dynamics and molecular mechanisms will become more popular, such as modeling cellular contractile forces from images of focal adhesion protein^45^.

Cell size is the most well studied aspect of shape in literature, it has been indicative of vast functions across species^46^. Cell size changes across the cell cycle, which can lead to altered protein concentrations depending on whether their synthesis or degradation scale proportionally^46–48^. Indeed, we found that the expression of certain proteins changes proportionally with cell and nuclear size, cell-nuclei-ratio, and cell cycle. Other than size and nuclear polarization, the width-height shapemode also provides insights and high associations with expression variability of many proteins, under normal and perturbed conditions. For example, we found differential metabolic profiles by shape-constriction/shape matched populations of large polarized cells that are thin or round in the G2 cell cycle phase, suggesting that they possibly prepare for different cell fates, as indicated by up/down regulation of proteins involved in different sets of metabolic pathways.

Intensity parameterization, whether through continuous warping or discrete spatial sampling, standardizes protein spatial representation to a common coordinate framework. The resulting spatial density distribution—at the protein or aggregated organelle level—provides a basis for comparing organellar topology. Furthermore, using the parameterized matrices to identify spatial latent factors allows us to observe the localization profiles of a protein in a cell population and to detect a shift from a heterogeneous protein distribution in non-perturbed cells to a more homogeneous localization under drug treatment. We leveraged this approach to identify proteins losing cytoplasmic concentration following vorinostat treatment. More broadly, parameterizing protein spatial distribution using a common coordinate framework serves as a powerful indicator of mechanism of actions following perturbation, since protein-relocalization is strongly associated with disease variants^49^.

This work provides an overall shapespace analysis across cell lines with proteome-wide measurement on the single-cell level. We define associations between shapes and organelles, pathways and spatial representation of individual proteins, and reveal that common coordinate frameworks anchored to cell and nucleus shape can serve as a unifying framework for comparative image-based studies of cells at subcellular resolution, which may include classic immunofluorescence microscopy, spatial proteomics, biosensor measurements, spatial transcriptomics, and future image-based technologies. For example, by aligning molecular measurement with morphological context, shapespace holds great promise for robust cross modality comparison with biological interpretability in perturbation assays such as optical pooled screening data, as it enables the detection of subtle, shape-constrained molecular changes (such as organelle association difference or protein localization shift) that are often masked in population-averaged analyses. Lastly, our study provides a foundation for proteome-wide exploration linked to shape variability, enabling the identification of key proteins for biophysical modeling to uncover the underlying principles governing physical cellular process.

## Methods

### Datasets

From the HPA subcellular section^50^, data from 10 cell lines in version 23 were included: osteosarcoma cell line U-2 OS, breast cancer cell line MCF7, epidermoid carcinoma cell line A431, glioblastoma cell line U251MG, kidney cancer cell line HEK293, liver cancer cell line HepG2; together with non cancer cell lines such as fibroblast cell line BJ, urinary bladder tissue cell line RT4, and two suspension cell lines HEL (erythroblast) and K562 (erythroleukemia) (Fig. 1A). The image data for each cell line includes between 5,242 to 262,802 individual cells, with proteins localizing to at least 10 organelles per cell line (Supplementary Table 1). This covers 11,998 protein-encoding genes.

The U2OS-FUCCI dataset^29^ was also derived from the subcellular section of the HPA. In this dataset of 357,083 cells from 1160 proteins displaying single cell heterogeneity were stained on U2OS-FUCCI cells and automatically imaged with a wide field microscope equipped with a 40x objective^29^. U2OS-FUCCI cells represent U-2 OS cells transgenically modified to tag two cell cycle regulators with fluorescent proteins: CDT1 (mKO2-hCdt1+), which accumulates in G1 phase, and GMNN (mAG-hGem+), which accumulates in S and G2 phases ^51^.

The perturbation dataset was obtained from the Cell Maps 4 Artificial Intelligence / Bridge2AI project ^52^. The specific imaging data set used in this study was published under the following persistent identifier: https://dataverse.lib.virginia.edu/dataset.xhtml?persistentId=doi:10.18130/V3/DXWOS5. The dataset contains images of the breast cancer cell line MDA-MB-468 in an untreated culturing condition or treated with the anti-cancer drugs paclitaxel or vorinostat. Each image features three cellular marker channels for nucleus, microtubules, and endoplasmic reticulum, and a channel for a protein of interest stained by one antibody. The data set contains 31,245 cells and stainings with 95 different antibodies to label a protein of interest.

### Cell and nuclear segmentation

#### HPACellSegmentator and single cell labels

For segmentation of single cells, a pre-trained DPN-UNet, which was the winning architecture in the 2018 Data Science Bowl^53^, was fine-tuned on HPA data. For each image, Nuclei, Microtubules and Endoplasmic Reticulum channels were used as a three-channel input and manually annotated cell masks were used as ground truth. Each channel had a frame size of 2048 x 2048 pixels and was acquired with a 63x objective with pixel resolution of 80 nm.

The DPN-UNet neural network was trained and validated on a dataset of 266 HPA Cell Atlas images from 28 cell lines, splitted for training (257 images) and validation (9 images). During inference, a pre-trained DPN-UNet for nuclei segmentation^53^ was used to generate nuclei mask images, followed by customized post-processing steps with watershed to obtain the individual nuclei. Then, using the DPN-Uet trained on our cell images, we obtained the cell masks. The nuclei obtained in the first step were combined with the cell mask in a watershed based post-processing step and produced the segmented cells. The activation function was softmax, and the loss function was a combination of Binary Cross Entropy (BCE) and Dice Loss. The model was trained with 1 Nvidia GeForce RTX 2070 until converged, which took roughly about 1.5 hrs (with pretrained weights).

The source code for the DPN-Unet segmentation model is available at https://github.com/cellProfiling/hpa-cell-segmentation.

DPNUnet was used to segment 61,463 FOV for 10 cell lines, resulting in 862,111 single cells after post processing removal of bordered cells. Each single cell label was predicted based on the output of an inceptionv3 model trained for single cell classification^28^.

#### Cellpose2.0

For segmentation of single cells in the U2OS-FUCCI dataset, we trained 2 small Cellpose2.0^54^ models for nuclei and cell segmentation. For the nucleus model, DAPI was used as 1-channel input. For each image, Nuclei, Microtubules and Endoplasmic Reticulum channels were used as 3-channel input and manually annotated cell masks were use

### Alignment of cell and nucleus orientation

Alignment of the cells was crucial to shape analysis. First, the cells and nucleus shapes were moved so that all nucleus centroids were centered to point (0,0) of a Catersian coordinate system. Cell and nuclei masks were aligned to the longest cell axis (general framework for all cell types) and cell centroid-to-nucleus centroid vector (for fixing polarity, as the majority of cell lines are of epithelial morphology). Since fourier transformation is sensitive to the orientation of shapes, each alignment resulted in a slightly different shapespace (i.e if the cells are aligned by cell-nucleus-centroid vectors, there’s no *‘polarity’* mode in shapespace).

### Shapemode identification

Treating contours as a wave upon itself, Fourier transformation decomposes the cell and nucleus outline into a harmonic series^55^. Specifically, from a point of alignment, each x,y coordinate vector was treated as a piecewise linear waveform tracing the cell shapes that can be reversibly transformed into the frequency domain (represented by 128 coefficients for eg). *numpy*^56^ implementation of Fast Fourier Transformation algorithm (*np.fft*) was used to calculate the complex fourier coefficients of nucleus *x*, nucleus *y*, cell *x*, cell *y* coordinates. The Hermitian symmetry allows reconstructing shapes based on half of the number of coefficients. These coordinates were joined into a single vector, making the whole shape datasets to be represented by a *(n_cells)* x *(4xFFTcoef)* matrix (since we used 128 fourier coefficients, the matrix size would be 264kx512 for U-2 OS and 254kx512 for U2-OS-FUCCI^29^ for examples). To reduce the dimensionality of the joint vectors of nucleus and cell coefficients, *scikit-learn*^57^ implementation of principal component analysis (PCA) and independent component analysis (FastICA) were used. 6 principal components (PCs) were found ∼90% of the variance (Figure 1A).

### Cell and nuclear shapespace construction

To identify what each PC represents, we z-scored all PCs independently by dividing the PC values by the standard deviation (σ) of that PC. The z-scored PCs were referred to as “shapemodes” and the combination of all 6 shapemodes made up the shapespace (Figure 1B). The usage of harmonic series as shape descriptors, followed by dimensionality reduction to construct shape variations were proposed in plant leaf analysis^58^ and recently used in 3D human cell shapes^7^.

### Spatially aware intensity parameterization through sampling and warping by deformation fields of cellular landmarks

After the primary modes of shape variation have been identified, we want to study how a single protein’s intensity or group of proteins’ intensity (*e.g.,* all proteins that perform X function or all proteins that make up Nucleoplasm) vary through different shapemodes. This requires mapping the protein signal into the shapemodes. In particular, protein intensity at certain locations could be sampled or warped into the average cell shape bin.

For sampling, 10 concentric rings from the nucleus centroid to nucleus membrane and similarly 20 concentric rings from nucleus membrane to the cell membrane were interpolated, resulting in 31 interpolated layers, each with 512 points. [10,20] was chosen because most cells in this study have epithelial morphology, hence the cytosolic space is large and dispersed compared to the nucleus. In each point, the raw protein signal was sampled at a 5×5 kernel centering at the interpolated point (Figure 1C). Each cell’s protein expression pattern is now represented as a numerical 31×512 matrix which was subsequently aggregated through each shapemode (Figure 1D). Before aggregation to represent organelle maps at specific shape bins, each matrix is binarized by image-level threshold. This method could suffer from oversampling and undersampling at certain regions for some cells (Supplementary Figure S6).

For warping, nucleus centroid, 32 points of nucleus contour and 64 points of cell contour were used as landmarks to morph protein expression of raw image to average cell. In specific, deformation fields of xx and yy were calculated from the landmark points of the average cell to those of each individual original cell, then based on these deformation fields, all pixel intensities can be reverse transformed/mapped back from original cell to average cell. Before aggregation to represent organelle maps at specific shape bins, each morphed cell representation is binarized.

The purpose of either of these intensity parameterization is to bring individual protein spatial expression in individual cells into a common reference framework. Sampling was used when matrix representation was beneficial, such as using matrices for latent factorization to get protein localization patterns in the breast cancer cell line for perturbation study where nucleus and cell area are relatively balanced. Warping with deformation fields, which create a smooth representation and mitigate under/oversampling problem, was used to generate average organelle maps across different organelles and cell lines, regardless of the organelle pattern and cytosol-nu-nucleus ratio.

### One- and two-dimensional distribution shift between pairs of average organelle representation

Replying on the fast closed form calculation of *scipy.stats.wasserstein_distance* for 1d wasserstein distance calculation, we can adopt the practical form of computing a Monte-Carlo approximation of the p-Sliced Wasserstein distance ^59^:

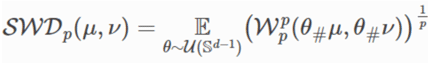

where:

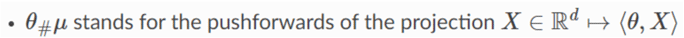

Consider each average organelle representation as the 2D (two-dimensional) spatial distribution of proteins that are made up of organelles.

In the space D^2, we can construct a probability distribution P(X_org) with the variable X_org. Here, by properly creating a set for space D^2, the probability distribution can be set for each population data in a desired specific region. The actual population (original sum pixel) can be obtained back by multiplying the probability (average org representation) by the total number of cells in that organelle population.

### Gene set enrichment analysis for variable proteins

1160 variable proteins stained in the U2OS-FUCCI cell line were enriched in MSigDB^60,61^ database against background 11,000 proteins expressed in U-2 OS from the HPA database. FDR < 0.05 was considered statistically significant. Each enriched term was visualized with a fraction of proteins variable with shapemode in a pie chart.

### Statistical testing

Pearson correlation, colocalization quotient, skewness tests in this study were performed with standard packages such as *numpy.corrcoef*, *scipy.stats.sknewtest*.

To assess the variability of protein expression in relation to shapespace, we applied binned and moving average statistics along each shapemode.

For binned analysis, we partitioned each shapemode into three groups: (1) the average shape group [−0.5\var,0.5\var] and two opposing groups corresponding to the lower [min,−0.5\var] and upper [0.5\var,max] extremes. For each protein, pairwise t-tests were conducted between the three groups (Group 1 vs. Group 2, Group 2 vs. Group 3, and Group 1 vs. Group 3). Additionally, a Kruskal-Wallis test was performed to assess overall differences across the three groups. To account for multiple testing, false discovery rate (FDR) correction was applied, and proteins with FDR < 0.05 in at least one of the three t-tests or the Kruskal-Wallis test were considered significant.

To assess continuous protein expression changes with shape, we examined whether protein expression follows a moving average trend along the shapemode projection. This approach quantifies how well protein expression is structured in relation to shape by calculating the explained variance. The percent variance explained by shapemode projection is defined as:

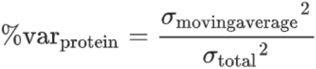

where σ_movingavg2 represents the variance of moving average values of protein expression ordered by shapemode position, and σ_total2 is the total variance of protein expression. A higher variance ratio indicates a stronger association between protein expression and shapemode variation. To evaluate significance, we performed permutation testing. The protein expression values were randomly shuffled across cells (PERMUTATIONS = 1000) to generate a null distribution of variance explained. The observed variance ratio was compared to the mean permuted variance ratio, and a significance threshold was determined by computing the difference between the two distributions.

### Latent factorization of intensity parameterization matrix

We applied non-negative matrix factorization (NMF) to the intensity parameterization matrices to identify the latent spatial components underlying the breast cancer dataset including untreated, vorinostat- and paclitaxel-treated cells. All matrices were decomposed into six non-negative components using the NMF class from *scikit-learn*. The factorization resulted in a sample-to-component matrix *W*, which was normalized to represent the spatial distribution of protein signal for each sample and can be used to assign each sample to a certain spatial cluster, and a component-to-feature matrix H, which after visualizing and reversed transformed into the average cell shape can provide interpretable spatial patterns, such as nuclear, sub cytoplasmic or cell periphery localisations, as shown in Figure 6D.

To assess the shift in spatial profiles of perturbed from non-perturbed cell populations for each protein, we quantified the spatial distributions using the Wasserstein distance. For example, spatial pattern distribution of perturbed cells, either the most dominating location pattern or a normalized profile, is compared to those of non-perturbed cells. Proteins were then ranked for each treatment based on the magnitude of this spatial profile shift (Supplementary Table 7).

## Code availability

Single cell labels were determined as described before^27^; the code is available at: https://github.com/CellProfiling/HPA_SingleCellClassification_Analysis. The code for constructing and analyzing the shapespace is available at: https://github.com/CellProfiling/2D_shapespace.

## Acknowledgements

We want to acknowledge the entire Human Protein Atlas team for data generation. Funding was provided to EL by the Wallenberg Foundation (2021.0346), Schmidt Futures, Göran Gustafsson Foundation, the Bridge2AI Program (NIH Common Fund; OT2 OD032742), the Cancer Cell Map Initiative (NCI Center for Cancer Systems Biology; U54 CA274502), Stanford Institute for Human Centered AI, and Param Hansa Philanthropies. TL was supported by Agilent Fellowship. JNH was supported by a Postdoctoral Fellowship from the Wenner-Gren Foundations and by an EMBO Postdoctoral Fellowship (ALTF 556-2022).

## Author Contributions

EL, TL, SR, MV designed the overall study; TL, MV, OW planned the 2D shapespace construction pipeline. TL implemented the code base for modeling, performed analysis and visualization. WL and EL contributed to the analysis. AC contributed to the cell cycle and metabolism investigations. JNH contributed to the perturbation investigations. TL, WL and EL wrote the manuscript with input from all authors.

## References

1. Hooke, R. Micrographia, or, Some Physiological Descriptions of Minute Bodies Made by Magnifying Glasses : With Observations and Inquiries Thereupon. (London : Printed by Jo. Marten, and Ja. Allestry, printers to the Royal Society, and are to be sold at their shop at the Bell in S. Paul’s Church-yard, 1665).

2. Gurcan, M. N. et al. Histopathological Image Analysis: A Review. IEEE Rev. Biomed. Eng. 2, 147–171 (2009).

3. Mousavikhamene, Z., Sykora, D. J., Mrksich, M. & Bagheri, N. Morphological features of single cells enable accurate automated classification of cancer from non-cancer cell lines. Sci. Rep. 11, 24375 (2021).

4. Bergert, M. et al. Cell Surface Mechanics Gate Embryonic Stem Cell Differentiation. Cell Stem Cell 28, 209–216.e4 (2021).

5. Nelson, C. M., Khauv, D., Bissell, M. J. & Radisky, D. C. Change in Cell Shape Is Required for Matrix Metalloproteinase-Induced Epithelial-Mesenchymal Transition of Mammary Epithelial Cells. J. Cell. Biochem. 105, 25–33 (2008).

6. Rangamani, P. et al. Decoding Information in Cell Shape. Cell 154, 1356–1369 (2013).

7. Viana, M. P. et al. Integrated intracellular organization and its variations in human iPS cells. Nature 613, 345–354 (2023).

8. Schaffer, L. V. et al. Multimodal cell maps as a foundation for structural and functional genomics. Nature 1–10 (2025) doi:10.1038/s41586-025-08878-3.

9. Théry, M. & Bornens, M. Cell shape and cell division. Curr. Opin. Cell Biol. 18, 648–657 (2006).

10. Minc, N., Burgess, D. & Chang, F. Influence of Cell Geometry on Division-Plane Positioning. Cell 144, 414–426 (2011).

11. Chen, T. et al. Large-scale curvature sensing by directional actin flow drives cellular migration mode switching. Nat. Phys. 15, 393–402 (2019).

12. Bodor, D. L., Pönisch, W., Endres, R. G. & Paluch, E. K. Of Cell Shapes and Motion: The Physical Basis of Animal Cell Migration. Dev. Cell 52, 550–562 (2020).

13. Rodrigues, M., Kosaric, N., Bonham, C. A. & Gurtner, G. C. Wound Healing: A Cellular Perspective. Physiol. Rev. 99, 665–706 (2019).

14. Lou, Y., Rupprecht, J.-F., Theis, S., Hiraiwa, T. & Saunders, T. Curvature-Induced Cell Rearrangements in Biological Tissues. Phys. Rev. Lett. 130, (2023).

15. Heisenberg, C.-P. & Bellaïche, Y. Forces in Tissue Morphogenesis and Patterning. Cell 153, 948–962 (2013).

16. Müllers, E., Silva Cascales, H., Jaiswal, H., Saurin, A. T. & Lindqvist, A. Nuclear translocation of Cyclin B1 marks the restriction point for terminal cell cycle exit in G2 phase. Cell Cycle Georget. Tex 13, 2733–2743 (2014).

17. Thul, P. J. et al. A subcellular map of the human proteome. Science 356, (2017).

18. Cho, N. H. et al. OpenCell: Endogenous tagging for the cartography of human cellular organization. Science 375, eabi6983 (2022).

19. Gut, G., Herrmann, M. D. & Pelkmans, L. Multiplexed protein maps link subcellular organization to cellular states. Science 361, eaar7042 (2018).

20. Black, S. et al. CODEX multiplexed tissue imaging with DNA-conjugated antibodies. Nat. Protoc. 16, 3802–3835 (2021).

21. Rood, J. E. et al. Toward a Common Coordinate Framework for the Human Body. Cell 179, 1455–1467 (2019).

22. Andersson, A., et al. A Landmark-Based Common Coordinate Framework for Spatial Transcriptomics Data. http://biorxiv.org/lookup/doi/10.1101/2021.11.11.468178 (2021) doi:10.1101/2021.11.11.468178.

23. Schaff, J. C. et al. SBML level 3 package: spatial processes, version 1, release 1. J. Integr. Bioinforma. 20, (2023).

24. Phillip, J. M., Han, K.-S., Chen, W.-C., Wirtz, D. & Wu, P.-H. A robust unsupervised machine-learning method to quantify the morphological heterogeneity of cells and nuclei. Nat. Protoc. 16, 754–774 (2021).

25. van Bavel, C., Thiels, W. & Jelier, R. Cell shape characterization, alignment, and comparison using FlowShape. Bioinformatics 39, btad383 (2023).

26. Schmied, C., Ebner, M., Ferré, P. S., Haucke, V. & Lehmann, M. OrgaMapper: A Robust and Easy-to-Use Workflow for Analyzing Organelle Positioning. http://biorxiv.org/lookup/doi/10.1101/2023.07.10.548452 (2023) doi:10.1101/2023.07.10.548452.

27. Janota, C. S. et al. Shielding of actin by the endoplasmic reticulum impacts nuclear positioning. Nat. Commun. 13, 2763 (2022).

28. Le, T. et al. Analysis of the Human Protein Atlas Weakly Supervised Single-Cell Classification competition. Nat. Methods 19, 1221–1229 (2022).

29. Mahdessian, D. et al. Spatiotemporal dissection of the cell cycle with single-cell proteogenomics. Nature 590, 649–654 (2021).

30. Zielke, N. & Edgar, B. A. FUCCI sensors: powerful new tools for analysis of cell proliferation. Wiley Interdiscip. Rev. Dev. Biol. 4, 469–487 (2015).

31. Liu, X., Jakubowski, M. & Hunt, J. L. KRAS gene mutation in colorectal cancer is correlated with increased proliferation and spontaneous apoptosis. Am. J. Clin. Pathol. 135, 245–252 (2011).

32. Ben-Sahra, I. & Manning, B. D. mTORC1 signaling and the metabolic control of cell growth. Curr. Opin. Cell Biol. 45, 72–82 (2017).

33. Kustatscher, G. et al. Understudied proteins: opportunities and challenges for functional proteomics. Nat. Methods 19, 774–779 (2022).

34. Chen, W.-C. et al. Functional interplay between the cell cycle and cell phenotypes. Integr. Biol. Quant. Biosci. Nano Macro 5, 523–534 (2013).

35. Kafri, R. et al. Cellular dynamics extracted from populations of fixed cells reveals a feedback between growth and progression through the cell cycle. Nature 494, 480–483 (2013).

36. Gut, G., Tadmor, M. D., Pe’er, D., Pelkmans, L. & Liberali, P. Trajectories of cell-cycle progression from fixed cell populations. Nat. Methods 12, 951–954 (2015).

37. Mou, P. K. et al. Aurora kinase A, a synthetic lethal target for precision cancer medicine. Exp. Mol. Med. 53, 835–847 (2021).

38. Magnusson, K. et al. ANLN is a prognostic biomarker independent of Ki-67 and essential for cell cycle progression in primary breast cancer. BMC Cancer 16, 904 (2016).

39. Pedley, A. M. & Benkovic, S. J. A New View into the Regulation of Purine Metabolism – The Purinosome. Trends Biochem. Sci. 42, 141–154 (2017).

40. Batta, G. et al. Alterations in the properties of the cell membrane due to glycosphingolipid accumulation in a model of Gaucher disease. Sci. Rep. 8, 157 (2018).

41. Alqosaibi, A. I. & Abdel-Ghany, S. Vorinostat induces G2/M cell cycle arrest in breast cancer cells via upregulation of PTEN. Eur. Rev. Med. Pharmacol. Sci. 27, 1503–1511 (2023).

42. Diz-Muñoz, A., Weiner, O. D. & Fletcher, D. A. In pursuit of the mechanics that shape cell surfaces. Nat. Phys. 14, 648–652 (2018).

43. Lecuit, T. & Lenne, P.-F. Cell surface mechanics and the control of cell shape, tissue patterns and morphogenesis. Nat. Rev. Mol. Cell Biol. 8, 633–644 (2007).

44. Bunne, C. et al. How to build the virtual cell with artificial intelligence: Priorities and opportunities. Cell 187, 7045–7063 (2024).

45. Schmitt, M. S. et al. Machine learning interpretable models of cell mechanics from protein images. Cell 187, 481–494.e24 (2024).

46. Fung, H. F. & Bergmann, D. C. Function follows form: How cell size is harnessed for developmental decisions. Eur. J. Cell Biol. 102, 151312 (2023).

47. D’Ario, M. et al. Cell size controlled in plants using DNA content as an internal scale. Science 372, 1176–1181 (2021).

48. Lanz, M. C., Fuentes Valenzuela, L., Elias, J. E. & Skotheim, J. M. Cell Size Contributes to Single-Cell Proteome Variation. J. Proteome Res. 22, 3773–3779 (2023).

49. Lacoste, J. et al. Pervasive mislocalization of pathogenic coding variants underlying human disorders. Cell 187, 6725–6741.e13 (2024).

50. Thul, P. J. et al. A subcellular map of the human proteome. Science 356, eaal3321 (2017).

51. Sakaue-Sawano, A. et al. Visualizing spatiotemporal dynamics of multicellular cell-cycle progression. Cell 132, 487–498 (2008).

52. Clark, T. et al. Cell Maps for Artificial Intelligence: AI-Ready Maps of Human Cell Architecture from Disease-Relevant Cell Lines. BioRxiv Prepr. Serv. Biol. 2024.05.21.589311 (2024) doi:10.1101/2024.05.21.589311.

53. Caicedo, J. C. et al. Nucleus segmentation across imaging experiments: the 2018 Data Science Bowl. Nat. Methods 16, 1247–1253 (2019).

54. Pachitariu, M. & Stringer, C. Cellpose 2.0: how to train your own model. Nat. Methods 19, 1634–1641 (2022).

55. Burger, W. & Burge, M. J. Fourier Shape Descriptors. in Principles of Digital Image Processing 169–227 (Springer London, London, 2013). doi:10.1007/978-1-84882-919-0_6.

56. Harris, C. R. et al. Array programming with NumPy. Nature 585, 357–362 (2020).

57. Pedregosa, F. et al. Scikit-learn: Machine Learning in Python. Mach. Learn. PYTHON 6.

58. Chitwood, D. H. & Sinha, N. R. Evolutionary and Environmental Forces Sculpting Leaf Development. Curr. Biol. 26, R297–R306 (2016).

59. Bonneel, N., Rabin, J., Peyré, G. & Pfister, H. Sliced and Radon Wasserstein Barycenters of Measures. J. Math. Imaging Vis. 51, 22–45 (2015).

60. Subramanian, A. et al. Gene set enrichment analysis: A knowledge-based approach for interpreting genome-wide expression profiles. Proc. Natl. Acad. Sci. 102, 15545–15550 (2005).

61. Liberzon, A. et al. The Molecular Signatures Database (MSigDB) hallmark gene set collection. Cell Syst. 1, 417–425 (2015).

