## Supplementary Figures for "Cell shapes decode molecular phenotypes in image-based spatial proteomics"

**Figure S1**: Cell line specific shapespace across 10 cell lines are similar.

**Figure S2:** Shapespace of U2OS-FUCCI for G1, G2S and G2 cell populations.

**Figure S3:** Similarity of the average cell line shape in common shapespace appears to align with the similarity in organelle correlation.

**Figure S4:** Association of meta data on PC1 (quantized).

**Figure S5:** Fluctuations in Minimum Cell Count Required for Robust Average Organelle Representation in U-2 OS Cells.

**Figure S6:** Gene set enrichment analysis of present genes for each cell line highlighting key biological pathways for that prospective cell line..

**Figure S7:** Intensity parameterization by sampling through concentric rings.

**Figure S8:** Average intensity from nuclear centroid of each organelle variation through shapespace.

### Supplementary Tables

***Table S1:*** Number of cells per organelle per cell lines from 10 cell lines in HPAv23.

***Table S2:*** PCA-identified shapemode percent variance for 10 cell lines.

***Table S3:*** Number of genes per cell line from 10 cell lines in HPAv23.

***Table S4:*** Gene set enrichment for Human Molecular Signatures Database.

***Table S5:*** Protein variation along each shapemodes with Kruskal-Wallis test, FDR < 0.05.

***Table S6:*** Protein variation along ICA-identified shapemodes with permutation test.

***Table S7:*** Protein localization patterns shift upon drug (vorinostat and paclitaxel) treatment.

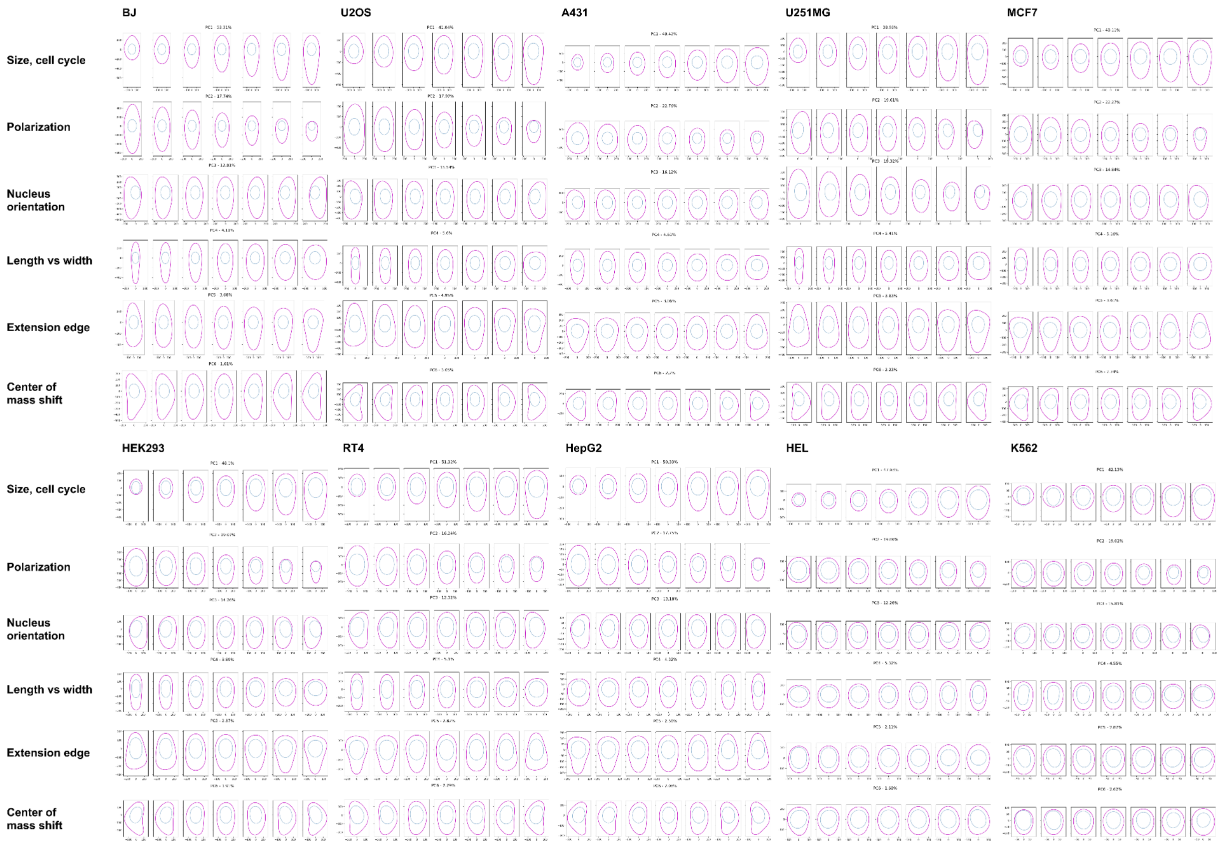


***Figure S1. Cell line specific shapespace across 10 cell lines are similar.***

*Cell line-specific shapespaces were constructed individually for osteosarcoma cell line U-2 OS, breast cancer cell line MCF7, epidermoid carcinoma cell line A431, glioblastoma cell line U251MG, kidney cancer cell line HEK293, liver cancer cell line HepG2, fibroblast cell line BJ, urinary bladder tissue cell line RT4, erythroblast cell lines HEL and erythroleukemia cell line K562. After applying fourier transformation to the contour coordinates of the cell and nucleus, the resulting shape coefficients were combined into a shape matrix, followed by principal component analysis (PCA) to capture major modes of variation. All shapespaces consistently exhibit six principal shapemodes, reflecting variation in size, polarization, nuclear orientation, length vs width, extension edge and center of mass shift.*

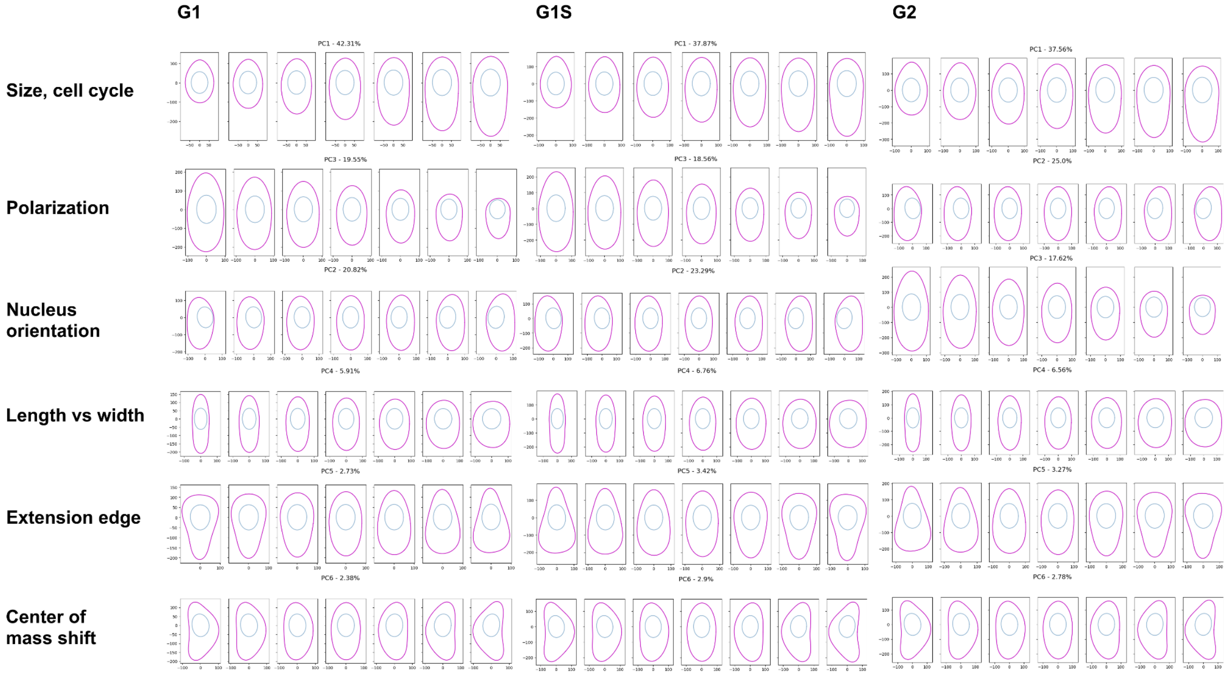


***Figure S2. Shapespace of U2OS-FUCCI for G1, G2S and G2 cell populations.***

*The shapespaces of U2OS-FUCCI cell populations in G1, G2/S, and G2 phases are also similar, indicating shared dominant modes of morphological variation across cell cycle states despite average cell size growth.*

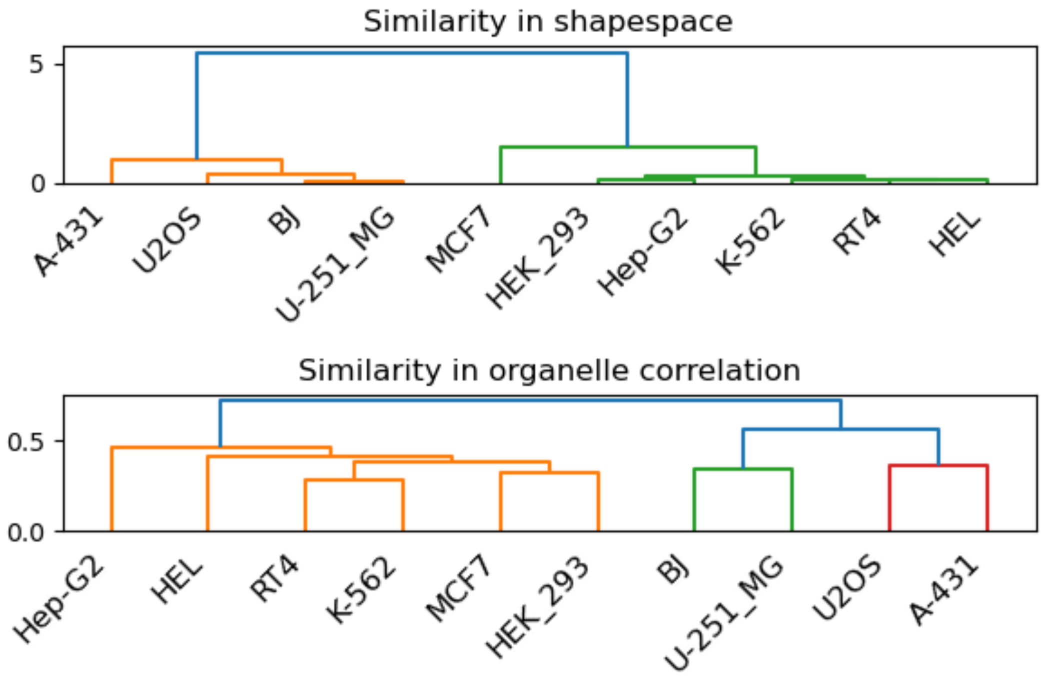


***Figure S3. Similarity of the average cell line shape in common shapespace appears to align with the similarity in organelle correlation.***

*Similarity in shapespace is calculated by the correlation of cell line shape across PCs in common shapespace. Similarity in organelle topology is measured by mean absolute distance between the pairwise organelle correlations. For each cell line pair, only organelle pairs present in both cell lines are taken into consideration. Clustering the similarity matrices in shape and topology space gives us two big clusters: BJ, A-431, U-2 OS, U-251 MG, and the rest of the cell lines.*

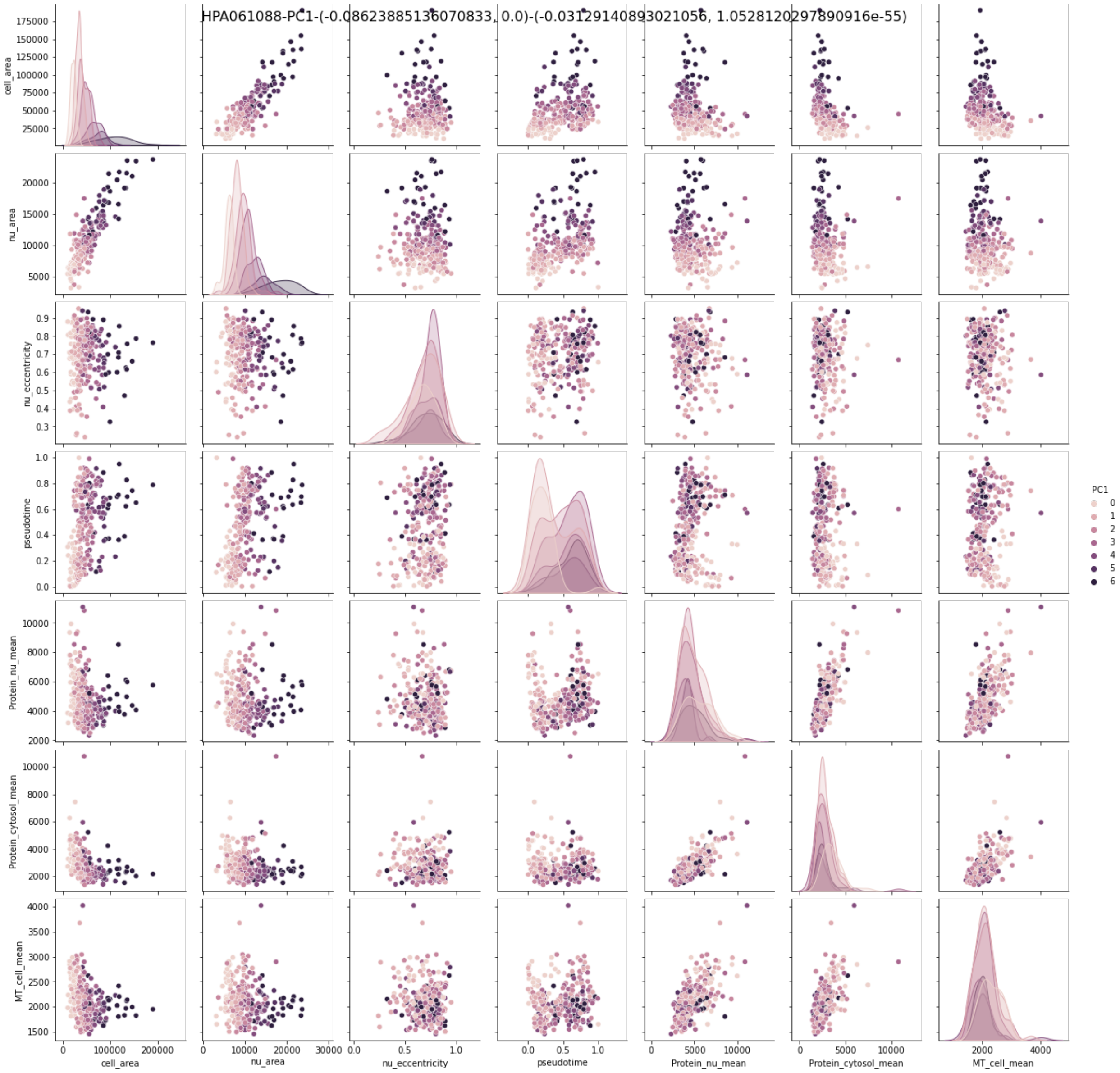


***Figure S4. Association of meta data on PC1 (quantized).***

*The first shapemode (PC1) of shape space, quantized into bins, is (partially) associated with multiple metadata features, including cell area, nuclear area, nuclear eccentricity, cell cycle pseudotime, average protein abundance in the nucleus and cytosol, and average microtubule abundance.*

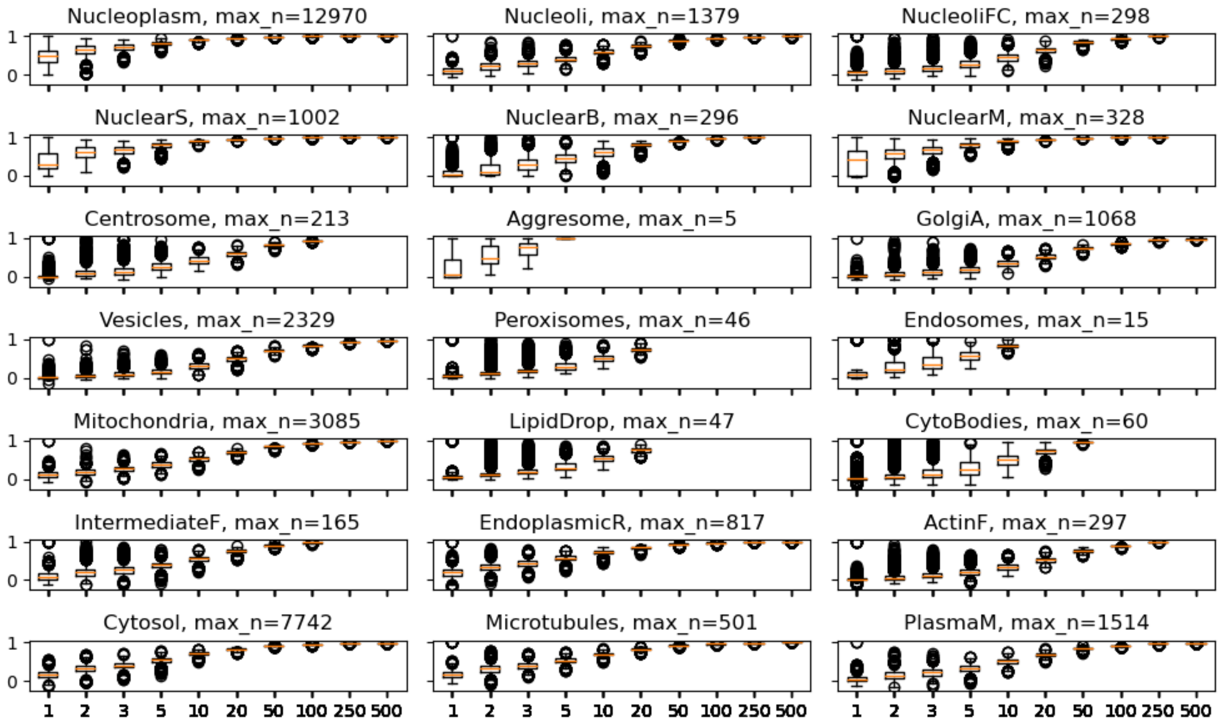


***Figure S5. Fluctuations in Minimum Cell Count Required for Robust Average Organelle Representation in U-2 OS Cells.***

*Stability of average organelle representation as a function of n-cell correlations. Most organelles reach a stable average representation with 50–100 cells, though the number required varies by organelle type. More stable organelles, such as the nucleoplasm and nuclear membrane, reach saturation with fewer cells, indicating lower variability across individual cells. In contrast, less stable (more dynamic) organelles, including intermediate filaments, plasma membrane, and actin filaments, require a higher cell count to achieve a robust average, reflecting greater morphological heterogeneity.*

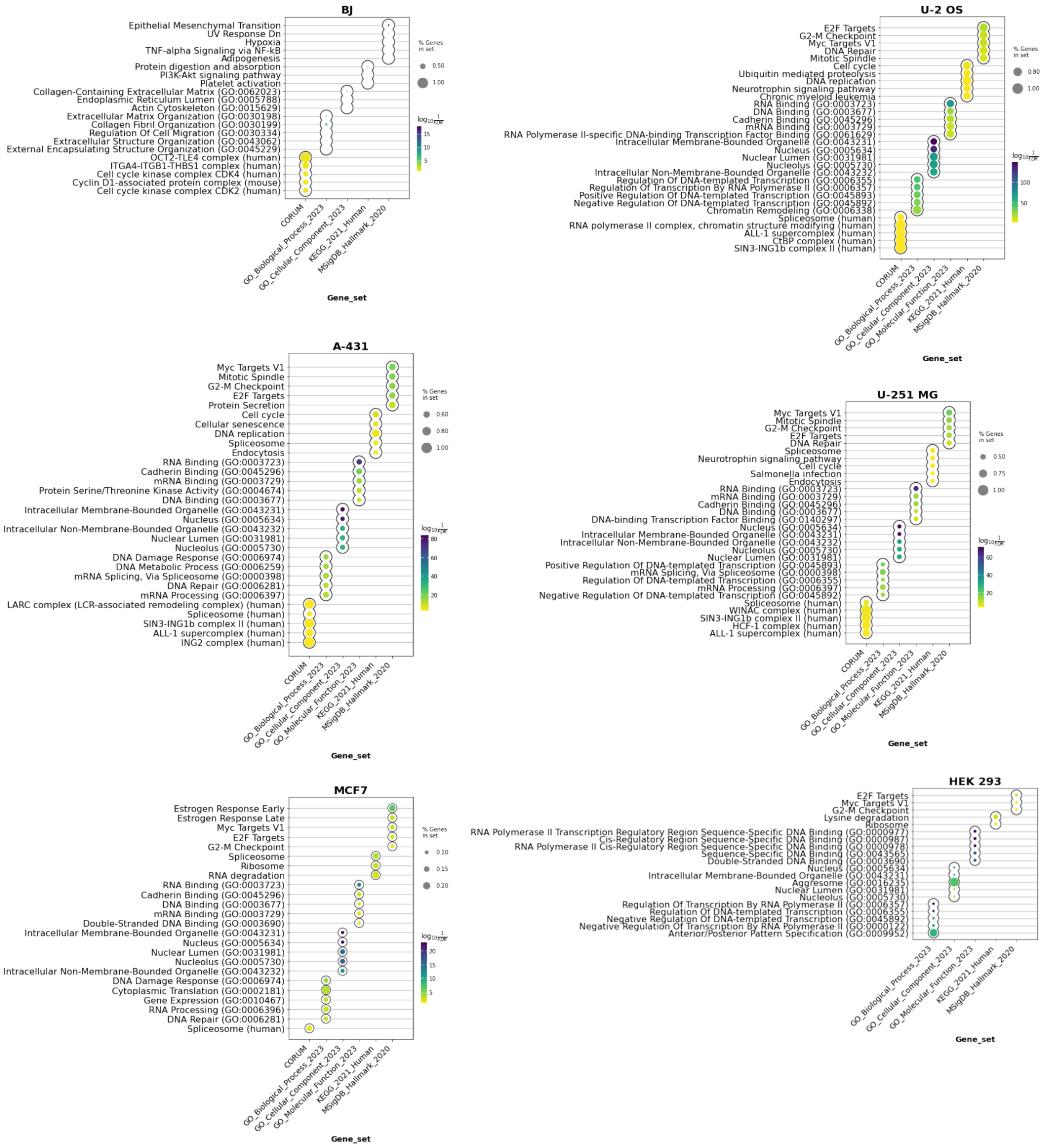
BJ
Epithelial Mesenchymal Transition
UV Response Dn
Hypoxia
TNF-alpha Signaling via NF-kB
Adipogenesis
Protein digestion and absorption
PI3K-Akt signaling pathway
Platelet activation
Collagen-Containing Extracellular Matrix (GO:0062023)
Endoplasmic Reticulum Lumen (GO:0005788)
Actin Cytoskeleton (GO:0015629)
Extracellular Matrix Organization (GO:0030198)
Collagen Fibril Organization (GO:0030199)
Regulation Of Cell Migration (GO:0030334)
Extracellular Structure Organization (GO:0043062)
External Encapsulating Structure Organization (GO:0045229)
OCT2-TLE4 complex (human)
ITGA4-ITGB1-THBS1 complex (human)
Cell cycle kinase complex CDK4 (human)
Cyclin D1-associated protein complex (mouse)
Cell cycle kinase complex CDK2 (human)
CORUM
GO_Biological_Process_2023
GO_Cellular_Component_2023
KEGG_2021_Human
MSigDB_Hallmark_2020
Gene_set
% Genes in set
0.50
1.00
15
10
5
U-2 OS
E2F Targets
G2-M Checkpoint
Myc Targets V1
DNA Repair
Mitotic Spindle
Cell cycle
Ubiquitin mediated proteolysis
DNA replication
Neurotrophin signaling pathway
Chronic myeloid leukemia
RNA Binding (GO:0003723)
DNA Binding (GO:0003677)
Cadherin Binding (GO:0045296)
mRNA Binding (GO:0003729)
RNA Polymerase II-specific DNA-binding Transcription Factor Binding (GO:0061629)
Intracellular Membrane-Bounded Organelle (GO:0043231)
Nucleus (GO:0005634)
Nuclear Lumen (GO:0031981)
Nucleolus (GO:0005730)
Intracellular Non-Membrane-Bounded Organelle (GO:0043232)
Regulation Of DNA-templated Transcription (GO:0006355)
Regulation Of Transcription By RNA Polymerase II (GO:0006357)
Positive Regulation Of DNA-templated Transcription (GO:0045893)
Negative Regulation Of DNA-templated Transcription (GO:0045892)
Chromatin Remodeling (GO:0006338)
Spliceosome (human)
RNA polymerase II complex, chromatin structure modifying (human)
ALL-1 supercomplex (human)
CtBP complex (human)
SIN3-ING1b complex II (human)
CORUM
GO_Biological_Process_2023
GO_Cellular_Component_2023
GO_Molecular_Function_2023
KEGG_2021_Human
MSigDB_Hallmark_2020
Gene_set
% Genes in set
0.80
1.00
100
50
A-431
Myc Targets V1
Mitotic Spindle
G2-M Checkpoint
E2F Targets
Protein Secretion
Cell cycle
Cellular senescence
DNA replication
Spliceosome
Endocytosis
RNA Binding (GO:0003723)
Cadherin Binding (GO:0045296)
mRNA Binding (GO:0003729)
Protein Serine/Threonine Kinase Activity (GO:0004674)
DNA Binding (GO:0003677)
Intracellular Membrane-Bounded Organelle (GO:0043231)
Nucleus (GO:0005634)
Intracellular Non-Membrane-Bounded Organelle (GO:0043232)
Nuclear Lumen (GO:0031981)
Nucleolus (GO:0005730)
DNA Damage Response (GO:0006974)
DNA Metabolic Process (GO:0006259)
mRNA Splicing, Via Spliceosome (GO:0000398)
DNA Repair (GO:0006281)
mRNA Processing (GO:0006397)
LARC complex (LCR-associated remodeling complex) (human)
Spliceosome (human)
SIN3-ING1b complex II (human)
ALL-1 supercomplex (human)
ING2 complex (human)
CORUM
GO_Biological_Process_2023
GO_Cellular_Component_2023
GO_Molecular_Function_2023
KEGG_2021_Human
MSigDB_Hallmark_2020
Gene_set
% Genes in set
0.60
0.80
1.00
80
60
40
20
U-251 MG
Myc Targets V1
Mitotic Spindle
G2-M Checkpoint
E2F Targets
DNA Repair
Spliceosome
Neurotrophin signaling pathway
Cell cycle
Salmonella infection
Endocytosis
RNA Binding (GO:0003723)
mRNA Binding (GO:0003729)
Cadherin Binding (GO:0045296)
DNA Binding (GO:0003677)
DNA-binding Transcription Factor Binding (GO:0140297)
Nucleus (GO:0005634)
Intracellular Membrane-Bounded Organelle (GO:0043231)
Intracellular Non-Membrane-Bounded Organelle (GO:0043232)
Nucleolus (GO:0005730)
Nuclear Lumen (GO:0031981)
Positive Regulation Of DNA-templated Transcription (GO:0045893)
mRNA Splicing, Via Spliceosome (GO:0000398)
Regulation Of DNA-templated Transcription (GO:0006355)
mRNA Processing (GO:0006397)
Negative Regulation Of DNA-templated Transcription (GO:0045892)
Spliceosome (human)
WINAC complex (human)
SIN3-ING1b complex II (human)
HCF-1 complex (human)
ALL-1 supercomplex (human)
CORUM
GO_Biological_Process_2023
GO_Cellular_Component_2023
GO_Molecular_Function_2023
KEGG_2021_Human
MSigDB_Hallmark_2020
Gene_set
% Genes in set
0.50
0.75
1.00
60
40
20
MCF7
Estrogen Response Early
Estrogen Response Late
Myc Targets V1
E2F Targets
G2-M Checkpoint
Spliceosome
Ribosome
RNA degradation
RNA Binding (GO:0003723)
Cadherin Binding (GO:0045296)
DNA Binding (GO:0003677)
mRNA Binding (GO:0003729)
Double-Stranded DNA Binding (GO:0003690)
Intracellular Membrane-Bounded Organelle (GO:0043231)
Nucleus (GO:0005634)
Nuclear Lumen (GO:0031981)
Nucleolus (GO:0005730)
Intracellular Non-Membrane-Bounded Organelle (GO:0043232)
DNA Damage Response (GO:0006974)
Cytoplasmic Translation (GO:0002181)
Gene Expression (GO:0010467)
RNA Processing (GO:0006396)
DNA Repair (GO:0006281)
Spliceosome (human)
CORUM
GO_Biological_Process_2023
GO_Cellular_Component_2023
GO_Molecular_Function_2023
KEGG_2021_Human
MSigDB_Hallmark_2020
Gene_set
% Genes in set
0.10
0.15
0.20
20
15
10
5
HEK 293
E2F Targets
Myc Targets V1
G2-M Checkpoint
Lysine degradation
Ribosome
RNA Polymerase II Transcription Regulatory Region Sequence-Specific DNA Binding (GO:0000977)
Cis-Regulatory Region Sequence-Specific DNA Binding (GO:0000987)
RNA Polymerase II Cis-Regulatory Region Sequence-Specific DNA Binding (GO:0000978)
Sequence-Specific DNA Binding (GO:0043565)
Double-Stranded DNA Binding (GO:0003690)
Nucleus (GO:0005634)
Intracellular Membrane-Bounded Organelle (GO:0043231)
Aggresome (GO:0016235)
Nuclear Lumen (GO:0031981)
Nucleolus (GO:0005730)
Regulation Of Transcription By RNA Polymerase II (GO:0006357)
Regulation Of DNA-templated Transcription (GO:0006355)
Negative Regulation Of DNA-templated Transcription (GO:0045892)
Negative Regulation Of Transcription By RNA Polymerase II (GO:0000122)
Anterior/Posterior Pattern Specification (GO:0009952)
GO_Biological_Process_2023
GO_Cellular_Component_2023
GO_Molecular_Function_2023
KEGG_2021_Human
MSigDB_Hallmark_2020
Gene_set
% Genes in set
0.10
0.20
0.30
15
10
5

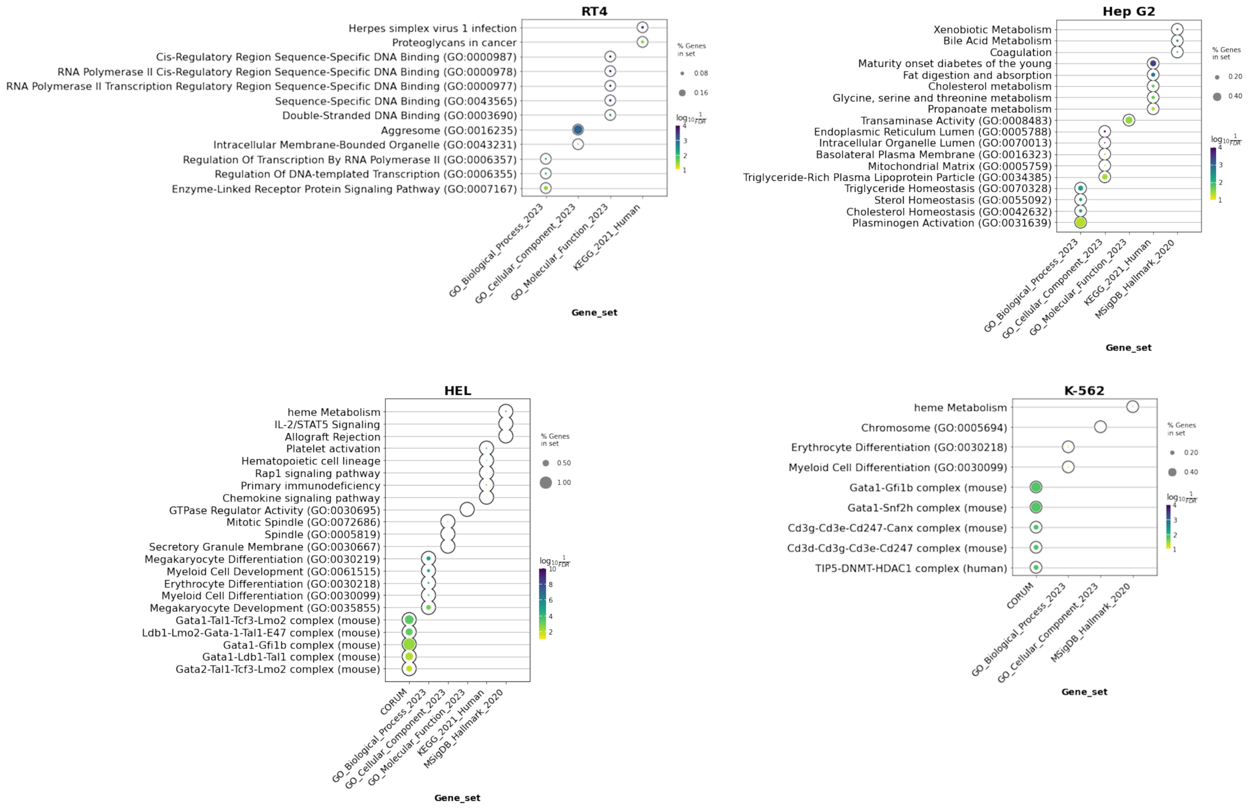


***Figure S6. Gene set enrichment analysis of present genes for each of the 10 cell lines highlighting key biological pathways for that prospective cell line.***

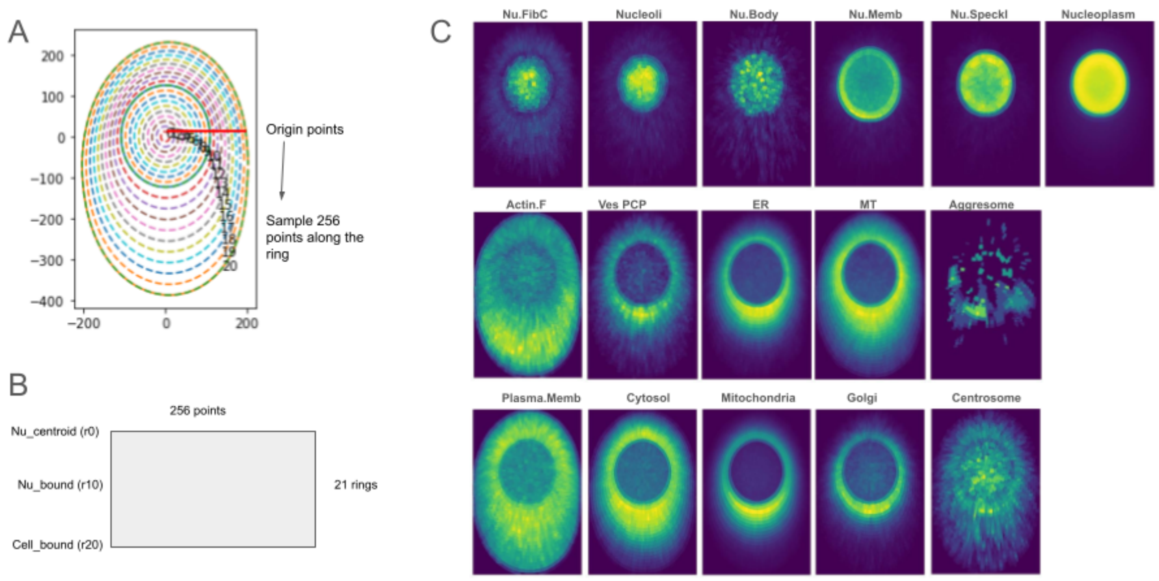


***Figure S7. Intensity parameterization by sampling through concentric rings.***

*(A) 10 concentric rings from nuclei centroid to nuclei membrane and 10 concentric rings from nuclei membrane to cell boundary were interpolated. Within each ring, 256 equidistant points are sampled. This gives rise to a square expression matrix (B), where nuclear proteins will have high expression in the upper part of the matrix, and cytosolic proteins will have high expression represented in the lower part of the matrix. The number of rings or sample points per ring can be adjusted based on experiments and cell type. However, to ensure the matrix is rectangular, the same number of points are sampled from each ring, leading to over representation in the nucleus and under representation in the cytosol, especially for cell lines with disproportionate nuclear-cytosol size. (C) Organelle average representation using matrix intensity parameterization, exemplified in the U-2 OS cell line.*

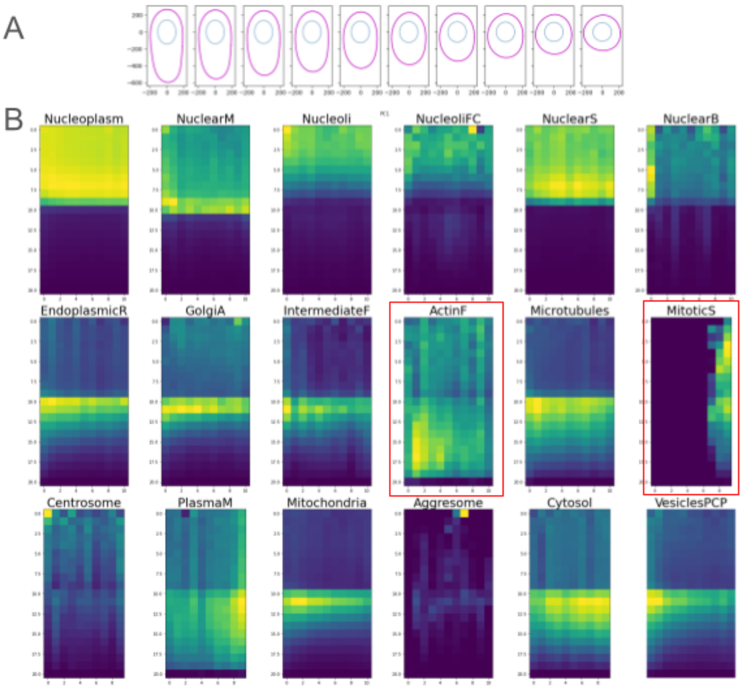


***Figure S8. Average intensity from nuclear centroid of each organelle variation through shapespace.***

*(A). PC1 of shapespace progression. (B) Variation of organelle representation as the cell size decreases. Most organelles have stable aggregated protein expression as cell shape changes, with a few exceptions. Proteins contributing to Mitotic Spindle only have presence and increased expression as the cell size decreases with respect to the nucleus size. Proteins contributing to Actin filaments have stronger expression when the cells are long and large, compared to round and small cells.*
